# Redox-enabled Electronic Interrogation and Feedback Control of Hierarchical and Networked Biological Systems

**DOI:** 10.1101/2023.08.22.554301

**Authors:** Sally Wang, Chen-Yu Chen, John R. Rzasa, Chen-Yu Tsao, Jinyang Li, Eric VanArsdale, Eunkyoung Kim, Fauziah Rahma Zakaria, Gregory F. Payne, William E. Bentley

## Abstract

We enable microelectronic devices to interrogate biology’s molecular communication, perform computations, and in real time control biological systems, including at several hierarchical levels: proteins, cells, and cell consortia. A key driver is establishing electronic access to and from biology’s native redox networks. First, redox-mediated electro-biofabrication facilitates facile assembly of biological components onto microelectronic systems, then electrode-actuated redox allows digital programming of enzyme activity, and further, redox-mediated electrogenetics facilitates closed-loop electronic control of cellular function. Specifically, we show algorithm-based feedback control of enzyme activity, cellular genetic circuits (via eCRISPR) and cell consortia behavior, all enabled by electronic data transfer. We further demonstrate electronic switching of cell-cell quorum sensing communication from one autoinducer network to another, creating an electronically controlled “bilingual” cell. We suggest these methodologies will not only help us to better understand biological systems, but design and control those currently unimagined.

---

Communication, or the transmission of information, builds the basis of the interconnected modern world we live in today. While electronic devices freely exchange information through electron transfer, devices that connect with biological processes are rare. We suggest this stems from the disparate communication modalities of biology and electronics^1,2^, their characteristic length and time scales^3^, and the orthogonal assembly processes of functioning units (cells vs. CPUs)^4^. Indeed, if one were to build an “Internet of Life” it would necessarily interact with and build on biology’s many communication modalities, including the transport of molecules, ions, and electrons, the latter being those that participate in redox reaction networks (**Fig. 1a**). It would be built of biofabricated structures, the designs for which naturally evolve, and are fabricated and dissimilated without impacting their surroundings. The emergence of transient electronics^5^ suggests our desire to create such a living information network, as our ability to eavesdrop on, compute, and take action on biological processes will be transformative for our society. Recognizing that redox-active molecules form the basis for electron transfer in biology, we suggest that redox can also serve to connect biological systems with electronics^1,2,6–10^. That redox-active molecules participate in redox reactions at electrode surfaces suggests that electrodes, in turn, can serve as a conduit for information exchange with biological systems, playing a pivotal role in a wide range of biological processes, including at the protein, cellular and multicellular levels.

**Figure 1.**
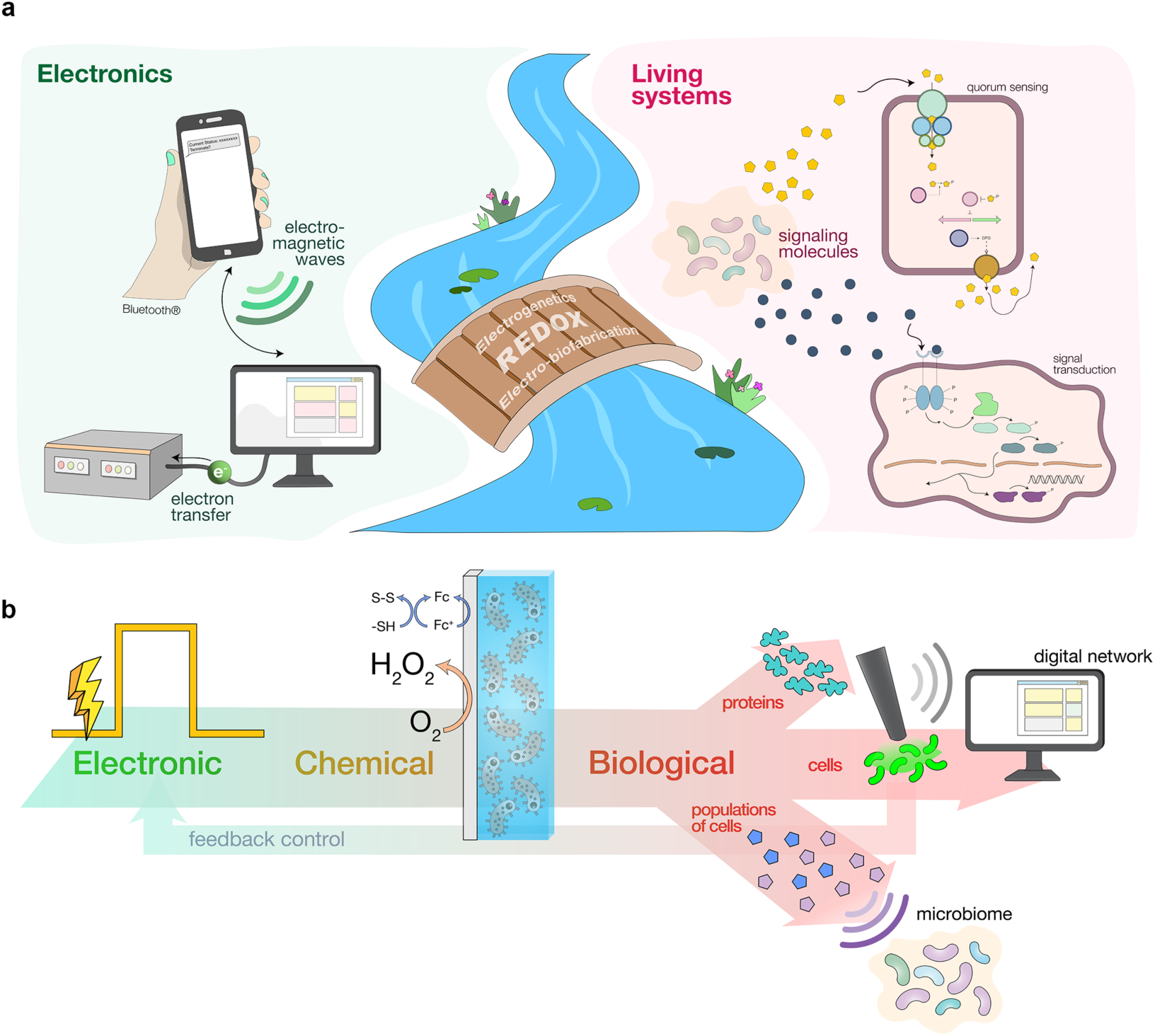
Bridging electronic and biomolecular communication through redox. **(a)** Electronic communication (left) mainly relies on free-flowing electron transfer or electromagnetic waves for communication. Molecular communication (right), conversely, employs signaling chemical molecules for information transfer. The redox modality can connect the two disparate communication modalities with redox-active molecules that can interact with both electronics and biology. Electro-biofabrication enables the creation of transmission interfaces and electrogenetics enables specific activation of engineered genetic circuits. **(b)** The redox signaling modality enables connection of biology to electronics in both directions. Here, an encoded electronic input is first transduced to chemical signals: (i) an oxidized redox mediator (Fc) that facilitates hydrogel assembly and (ii) reduced oxygen (H_2_O_2_) that is interpretable by several biological subsystems at the protein (top), cellular (middle), and multicellular (bottom) levels. Optical signals (e.g., fluorescence) generated by cells, as well as electrochemical currents (from peroxide generation), are recorded, computed upon, and fed back into process control algorithms for control, establishing an electro-bio-electro communication loop.

In this report, we develop methods for nearly seamless information transfer between the electronics of devices and the redox-based electronics of biology. This is enabled by electronic assembly of suitable interfaces, positioning of redox-active enzymes in configurations that provide rapid information transfer, as well as specific engineering of cells that enable precisely targeted transfer of electronic information to and from electrodes. Interfaces are assembled using redox chemistry mediated by electron carriers (e.g., ferrocene) in a way that preserves stability and viability for subsequent communication with electronics via the same electrode surface. We further show the creation of CRISPR^11,12^ genetic circuits that translate electronic information into biological forms, including the creation of an electronic “language translator” shifting a population of bacterial cells to operate in a bilingual manner. We then demonstrate how electroassembled enzymes and cells can provide in real time, optoelectronic forms of biological activity that can be computed on and used for closed loop control (**Fig. 1b**). Specifically, we show how electroassembled horseradish peroxidase (HRP) activity can be controlled on chip by the electronic synthesis of its substrate, hydrogen peroxide (henceforth, peroxide). We employ electrogenetics and the *oxyRS* stress regulon^13^ of *Escherichia coli* to demonstrate how engineered and native cells can be electronically stimulated (again through peroxide production) and recorded (via real time fluorescence and electrochemical measurements) for feedback electronic control of gene expression (**Fig. 1b**). Finally, in a manner analogous to the multifaceted and interwoven nature of the Internet of Life, we show how locally regulated biological systems can be networked and even controlled via a cascaded control architecture that includes a human interface (text messaging) that supervises model-based control actions and approves their implementation.

## Electro-biofabrication: assembly of biological components onto electrodes

Biofabrication^14^ has allowed biological components to be assembled on micro-, or nano-scaled electronic devices for biosensing^15–17^ or even control of biological functions^18^. Here, we enlisted electro-biofabrication^19^ of biocompatible hydrogels to facilitate the assembly of enzymes and cells onto electrodes to localize signal transfer. For enzyme assembly, we show the spatially-configurable electrodeposition of a covalently-conjugated gelatin/horseradish peroxidase (HRP)^20^ hydrogel film (**Figure 2a**, **Supplementary Fig. 1**). Its assembly builds on anodic oxidation^20^ but is fabricated here with spatially confined geometries enabling area-based interrogation and control of activity (**Figure 2b (ii)**)^21^.

**Figure 2.**
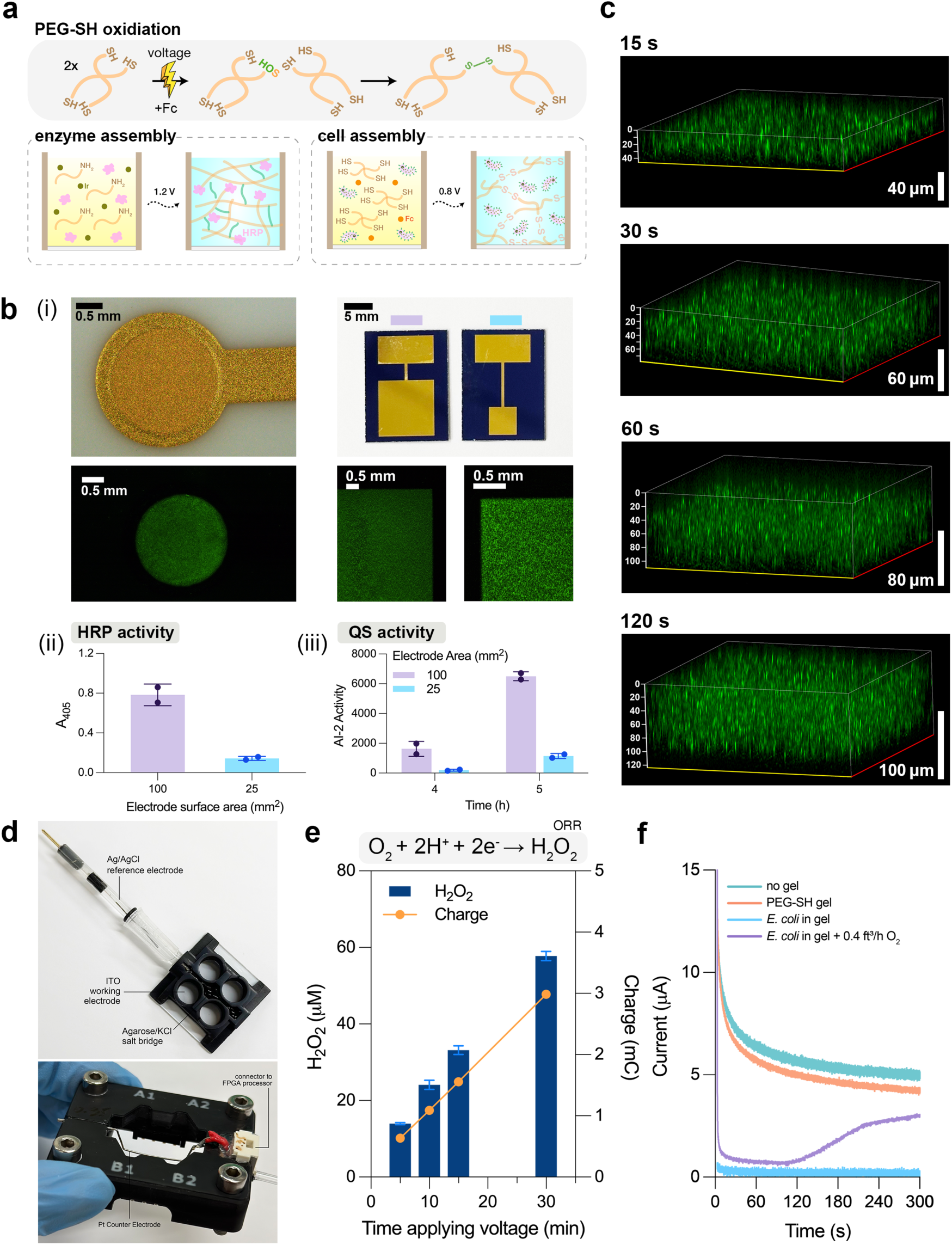
Electro-biofabrication for assembly of biological components and integration with an optoelectrochemical electronic system. **(a)** Schematic of PEG-SH oxidative crosslinking, assembly of HRP-conjugated gelatin hydrogel (enzyme assembly), and co-deposition of *E. coli* and PEG-SH (cell assembly). Redox mediators, Ir and Fc, facilitates the oxidation and subsequent gelation of gelatin and PEG-SH. **(b)** Spatially-programmable deposition of HRP/gelatin and *E. coli*/PEG-SH hydrogels. (i) Fluorescence microscopy images of “artificial biofilms” containing SYTO-9 dyed *E. coli* BL21 assembled onto circular 2 mm-diameter, square 5 × 5 mm, and square 10 × 10 mm gold electrodes. (ii) Activity of electroassembled HRP, represented as the absorbance at 405 nm through ABTS assay. (iii) Secreted AI-2 activity from electroassembled ‘artificial biofilms’ containing *E. coli* BL21 cells (OD_600_ = 5). Bar height represents the mean, filled circles represent individual replicates, and error bars represent the standard deviation (*n* = 2). **(c)** Representative Z-stack confocal images of the ‘artificial biofilms’ containing engineered *E. coli* that constitutively express GFP (DH5α-sfGFP). Deposition time is labeled on top of each image. X-axis is shown in yellow, and Y-axis is shown in red. Z-axis and scale bars are shown in white, all units are in μm. **(d)** ITO-based, 3D-printed optoelectrochemical device. The clear ITO glass serves as the working electrode and allows for optical observations. An Ag/AgCl electrode is used as the reference electrode, and a Pt wire (below, mounted on custom-fabricated connector) is the counter electrode. The three electrodes are connected through the salt bridge (1 M KCl in 1% agarose) casted in the central well of the device. **(e)** Peroxide (H_2_O_2_) generated (blue bar) in the ITO-based electrochemical platform as correlated to applied charge (orange circle). Bar height represents the mean, and the error bars represent the standard deviation (*n* = 4). **(f)** Current obtained during peroxide generation. Working electrode (WE) was poised at −0.8 V for 300 s for all samples. Teal: 150 μl of 20% LB. Orange: 150 μl of 20% LB submerging a cell-free PEG-SH film deposited on WE. Blue: 150 μl of 20% LB submerging an “artificial biofilm” containing OD_600_ = 6 of *E. coli* after 1.5 h of incubation at 34°C. Purple: 0.4 ft^3^/h of O_2_ supplied (starting at 0 s, through built-in tubing in the connector) to 150 μl of 20% LB submerging an “artificial biofilm” containing OD_600_ = 6 of *E. coli* after 1.5 h of incubation at 34°C.

For cell assembly, we took inspiration from bacterial biofilms found in nature and created ‘artificial biofilms’, consisting of *E. coli* entrapped in a thiolated polyethylene glycol (PEG-SH) hydrogel. Cells were mixed with a solution containing 4-armed PEG-SH monomer (50 mM) and redox mediator, ferrocene (Fc, 5 mM), to facilitate the electrode-induced oxidation of thiol groups into disulfide bonds, creating a crosslinked hydrogel^20^. When an oxidative voltage (slight variations depending on electrode material) is applied, PEG-SH crosslinks, forming a film that entraps cells within the growing matrix (**Fig. 2a**). In **Fig. 2b (i)**, PEG-SH hydrogel containing SYTO 9-stained *E. coli* BL21 (OD_600_ ∼ 5) are electro-assembled onto on patterned gold electrodes (+0.7 V, 2 min) with various geometries and surface areas (ranging from ∼13 mm^2^ to 100 mm^2^). The cell distribution appears homogeneous and clearly defined by the geometry of the electrode. 4-6 h after *E. coli* BL21 that constitutively secrete quorum sensing (QS) autoinducer AI-2 were assembled onto electrodes with different surface areas (25 mm^2^ vs 100 mm^2^), the growth media submerging the larger electrode contained ∼6 times the AI-2 activity of the smaller electrode, suggesting preservation and proportionality of cell function with electrode area (**Fig. 2b (ii)**). In **Fig. 2c**, ‘artificial biofilms’ are controllably deposited with custom thickness by varying the deposition time. Here, we deposited the films containing GFP-expressing DH5α-sfGFP on an optically-transparent indium tin oxide (ITO) electrode and quantified confocal Z-stack images to estimate their thickness. Results show the film thickness was correlated with deposition time, thus providing an additional orthogonal approach for designing and controlling characteristics of this “living” material.

To better facilitate optical assessment of electroassembled materials and cells, we transitioned to optically clear conductive indium tin oxide (ITO) electrodes. In **Fig. 2d**, we 3D-printed a four-well optoelectrochemical device with an ITO-coated working electrodes (WE) in a prototypical three-electrode setup. Here, each well shares the same Ag/AgCl reference electrode and Pt wire counter electrode through the salt bridge casted in the central well (**Supplementary Methods**). This optoelectrochemical device provides for electronic I/O, where the same electrode for assembly is used for electrochemical detection as well as electrochemical reduction of oxygen (oxygen reduction reaction (ORR); O_2_ + 2H^+^ + 2e^−^ ↔ H_2_O_2_) forming a transmitted redox signal, peroxide^7,22^. In **Fig. 2e**, we found that biasing the ITO electrode with −0.8 V produced up to ∼60 μM of peroxide (in 30 min), exhibiting linear dependence with duration and consequently, charge (product of time and current). We then investigated peroxide generation in the presence of a metabolizing ‘artificial biofilm’ in **Fig. 2f**. Measured current at an applied potential reflects the extent of the ORR reaction^23^. We observed dramatically reduced current when the PEG-SH films contained respiring *E. coli*, suggesting an oxygen limitation at the electrode surface for peroxide generation. That is, with the identically assembled biofilm, we supplemented 0.4 ft^3^/h of oxygen through a built-in gas transport tube (**Supplementary Fig. 10**), and an increased current was observed. In addition to demonstrating the importance of oxygen transfer to the electrode surface, this result also shows how the electrode of assembly can serve as a dynamic electrochemical sensor of the redox reaction and perhaps indirectly, the respiratory activity of the assembled cells.

Together, these results show the electro-assembly of redox-active hydrogels (gelatin and PEG-SH) can be configured with various geometries (area, thickness). Their biocompatibility also allowed both materials to either conjugate with enzymes or entrap live cells while retaining the biological component’s activity. Altogether, we built a biohybrid, optoelectrochemical device consisting of a living ‘artificial biofilm’ and an optically clear electronic platform capable of generating peroxide as the redox signal.

## oxyRS-based electrogenetic CRISPR activation (CRISPRa)

We then enlisted the versatile CRISPR transcriptional regulation system^11^ to allow activation, repression, and multiplexed transcriptional regulation of genes controlled by *E. coli’s* global oxidative stress regulon, *oxyRS*. The single guide RNA (sgRNA) was placed downstream of the *oxyS* promoter (**Fig. 3a**), and we monitored CRISPRa activity by the expression of *gfpmut2*. First, we confirmed in suspension cultures that both CRISPR sgRNA transcription and GFP expression were peroxide-inducible (**Supplementary Fig. 2a, b**). Next, we applied reducing potential (−0.8 V) to electroinduce CRISPRa in the cells electrically assembled on the ITO electrode surface (**Fig. 3b**). Using confocal microscopy, we found that ∼70 % of electroinduced cells expressed GFP, while ∼35% of electroassembled cells expressed GFP when 200 μM peroxide was introduced to the growth media above the film, and < 5% of the cells expressed GFP in negative controls. Interestingly, the fraction of fluorescing cells increased with charge (**Fig. 3c-d**).

**Figure 3.**
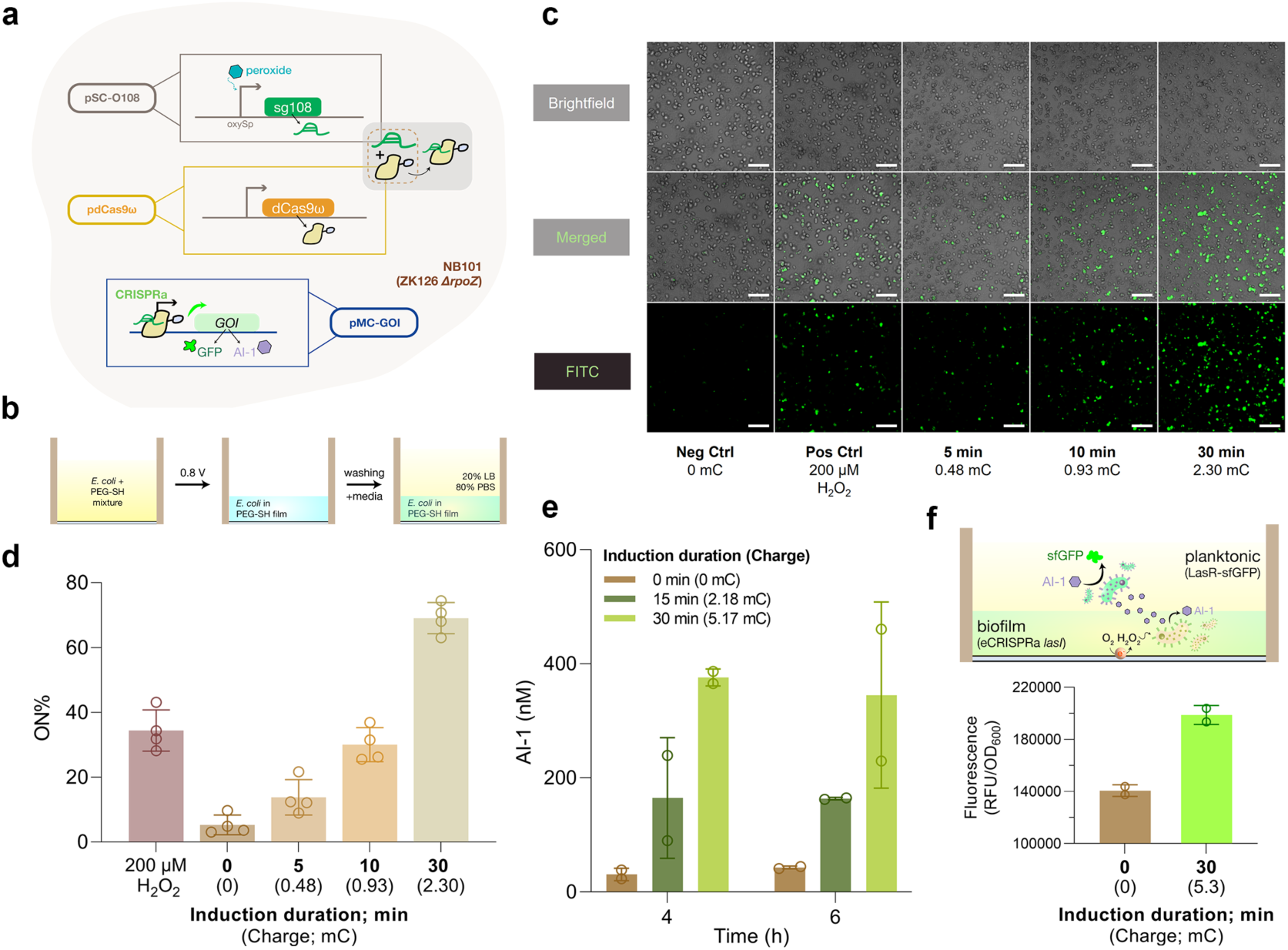
Tunable eCRISPRa within an “artificial biofilm”. **(a)** Schematic illustration of the electrogenetic CRISPRa system powered by the *oxyRS* regulon. Electro-induced gRNA sg108 (from plasmid pSC-O108) forms a complex with constitutively-expressed dCas9ω to initiate transcription of the gene-of-interest (GOI) in plasmids pMC-GFPmut2 or pMC-LasI-LAA. **(b)** General workflow and setup for experiments with ‘artificial biofilms’. After electrodeposition, we thoroughly washed the film with PBS to remove excess cell/PEG-SH solution. 150 μL of 20% LB was then added into the well to submerge the film in growth media. **(c)** Confocal images of *E. coli* harboring the eCRISPRa-GFP genetic cassette that were embedded in the PEG-SH film. Varying times of electroinduction and the resulting charges (millicoulomb; mC) applied to the cells are indicated below images. 200 μM of H_2_O_2_ was exogenously added to culture media to activate the *oxyS* promoter as the positive control. Top: brightfield; Middle: Merged; Bottom: FITC filter. **(d)** Percentage of *E. coli* in the PEG-SH film that were activated through eCRISPRa. Data is quantified from images in Fig. 3c with ImageJ. **(e)** AI-1 assay indicating the amounts of AI-1 generated via CRISPRa *lasI* cells. **(f)** Measured fluorescence in a coculture comprised of eCRISPRa *lasI* cells (in film) and AI-1 fluorescent reporters (NEB10β + LasR_S129T-GFPmut3) situated in the liquid above. For all bar graphs, the bar height represents the mean, and error bars represent the standard deviation (*n* ≥ 2). Individual replicates were indicated as open circles.

Having shown that we can electrically induce *oxyRS*-based CRISPRa in the optoelectrochemical device, we then sought to demonstrate information routing between electronic to biological signaling modalities by initiating a specific QS communication foreign to *E. coli* (i.e., *las* “AI-1” QS system from *Pseudomonas aeruginosa*) through peroxide-mediated CRISPR. A new reporter plasmid (pMC-lasI-LAA) was constructed by replacing *gfpmut2* with the AI-1 producer *lasI* (fused with an ssRA tag), allowing CRISPRa control of AI-1 production. Prior to electroinduction tests, CRISPRa *lasI* cells were confirmed to be peroxide-inducible in liquid culture (**Supplementary Fig. 3a**). Similar to the results of eCRISPRa-induced GFP which routes signals based on *soxRS* and supplemented redox mediators^11^, we found that AI-1 levels were directly electrically inducible via H_2_O_2_/*oxyRS* regulon and correlated to the duration of voltage application and charge (**Fig. 3e**). We then further demonstrated message transfer to a second population through CRISPRa-mediated QS signal routing in a consortia setup. Specifically, QS signal (AI-1) secreted from the electrode-bound CRISPRa *lasI* cells is relayed to the planktonic AI-1 responsive population (NEB10β harboring plasmid pLasR_S129T-GFPmut3, **Supplementary Fig. 3c**). After receiving electroinduction, the distant planktonic AI-1 reporters exhibited ∼1.5-fold increase in GFP levels compared to those in the non-induced co-culture (**Fig. 3f**).

In summary, we demonstrated on-device, peroxide-mediated eCRISPR that allows propagation of the localized electrogenetic cue across different cell populations through the electroassembled hydrogel film, and subsequent coordination of population-wide behavior with the help of QS communication. It is important to note that because we assembled cells via entrapment in a crosslinked hydrogel film, there was no need to engineer assembly features into the assembled cells^7^. There was also no need to supplement with message-conveying redox mediators, since the signal was directly transferred from the electrode. Thus, we anticipate virtually any cell or consortia of cells can be assembled and electronically stimulated using this redox-based or other stimulus-responsive, electroassembly methodologies^19,24^.

## eCRISPR inhibition and multiplexed control of QS to enable ‘bilingual’ communication

While we showed that enabling peroxide-mediated eCRISPR activation of QS can open a new line of communication to other microbial populations, CRISPR transcription regulation offers multiplexed function, including coincident activation and inhibition^25^. Here, we explored multiplexed control for multi-locus transcriptional regulation to enable electrically-controlled ‘bilingual’ QS communication. In particular, we aimed to create an engineered *E. coli* that switches its ‘spoken language’ (i.e., QS signaling) from its ‘mother tongue’ (*luxS*-based AI-2) to a ‘foreign language’ (*las* AI-1 from *P. aeruginosa*) upon receiving the electronic signal. Unlike the *las* system, *luxS/*AI-2 QS system is found natively in many bacteria, including our experiment chassis *E. coli*^26–28^. To prohibit *E. coli* from secreting AI-2, we designed the sgRNA LuxS1 that is homologous to the AI-2 producer gene *luxS* and repurposed dCas9ω for CRISPR inhibition (CRISPRi) (**Fig. 4a**). The gRNA, along with a non-specific control, was expressed initially under a strong, constitutive promoter in suspension culture to examine the efficacy of LuxS1. Compared to the control gRNA, we observed a 372-fold decrease in AI-2 activity from the cultures that constitutively expressed LuxS1, demonstrating highly effective inhibition (**Supplementary Fig. 4a**). Next, we replaced the constitutive promoter with the *oxyS* promoter to allow inducible expression of LuxS1. We found that AI-2 activity of the non-induced control was ∼7-fold higher than that of the induced sample, confirming peroxide-inducible inhibition of AI-2 QS system via CRISPRi (**Supplementary Fig. 4b**). Prior to testing electrogenetic CRISPRi, AI-2 profiles of in-film, electrode-bound cells were shown to share a similar (increasing) yet delayed trajectory with that of the suspension cultures^29^ (**Fig. 2**, **Supplementary Fig. 4c**). eCRISPRi of *luxS* was subsequently found; as shown in **Fig. 4b**, on-electrode CRISPRi cells that were subjected to electroinduction consistently exhibited lower AI-2 activity compared to the uninduced control, proving controllable quenching of AI-2 QS through eCRISPR.

**Figure 4.**
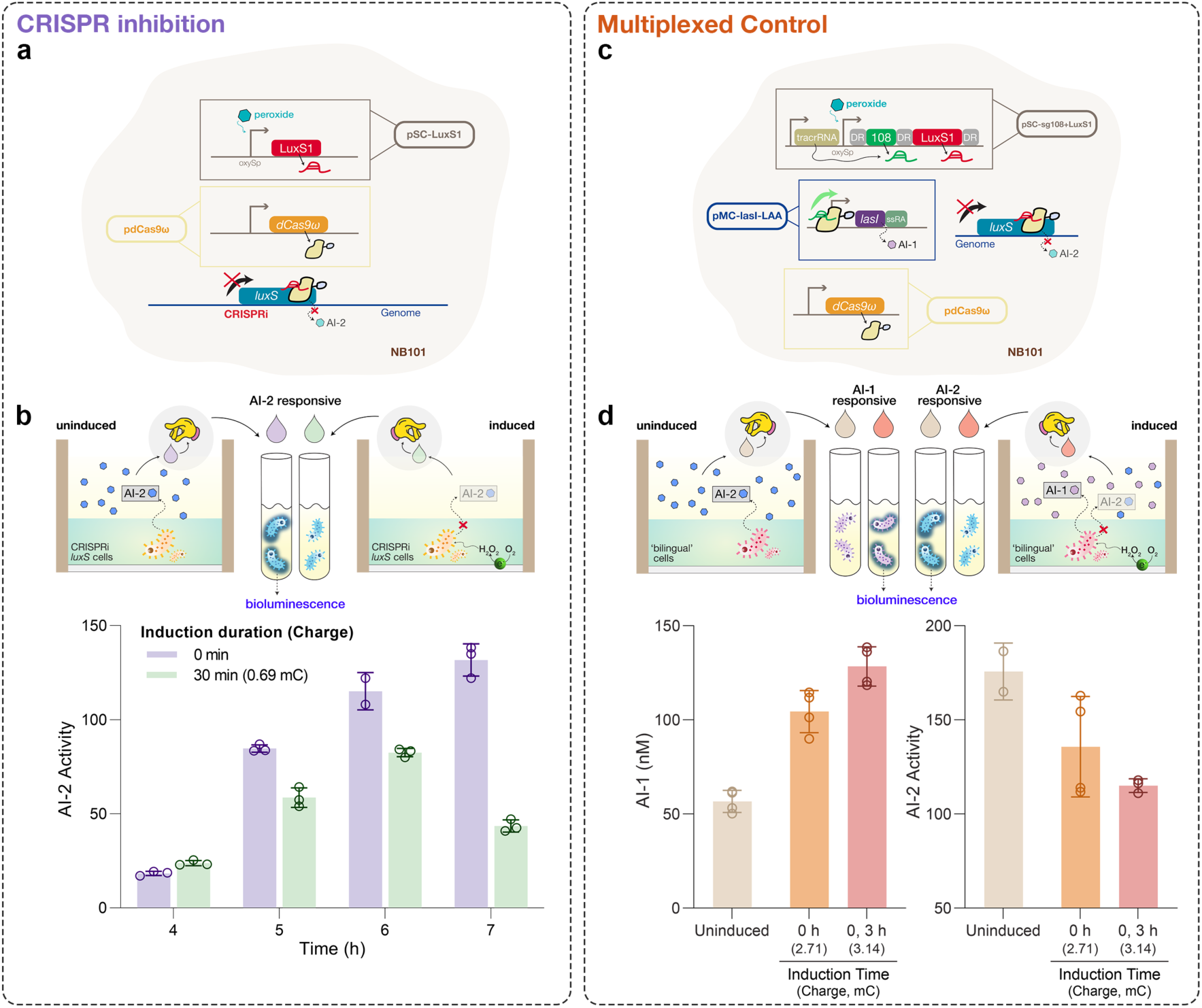
eCRISPR inhibition of a native QS signal and multiplexed control of QS communication. **(a)** Schematic of *luxS* eCRISPRi to inhibit AI-2 QS signaling. *luxS* specific gRNA LuxS1 is expressed under *oxyS* promoter in pSC-LuxS1. Both pSC-LuxS1 and pdCas9ω were transformed into NB101 allowing expression of gRNA and dCas9ω. **(b)** Measured AI-2 activity of the filtered media samples collected at various times post-induction. In-film eCRISPRi cells were electro-induced for 0 or 30 minutes 3 h after deposition. **(c)** Schematic of multiplexed eCRISPR for engineering a ‘bilingual’ strain. gRNAs sg108 and LuxS1 were flanked with the DR sequence and placed downstream of the *oxyS* promoter in pSC-sg108+LuxS1. tracrRNA was individually expressed under a synthetic constitutive promoter. Both pSC-LuxS1 and pdCas9ω were transformed into NB101 allowing gRNA expression. Plasmids pSC-sg108+LuxS1, pdCas9ω, and pMC-lasI-LAA were all transformed into NB101 allowing multiplexed gRNA expression. **(d)** Measured AI-1 levels and AI-2 activity of the filtered media samples collected 7 h post-deposition. In-film ‘bilingual’ cells were electro-induced for 0 or 30 minutes at 0 or 0 and 3 h post-deposition. For all figures, the bar height represents the mean, and the error bars represent the standard deviation (*n* ≥ 3). Individual replicates are indicated as open circles.

Having shown successful eCRISPRi of *luxS*, we then sought to harness the multiplexed nature of CRISPR and combine CRISPRa of *lasI* and CRISPRi of *luxS* to create a ‘bilingual’ strain. Though many strategies for multiplexed gRNA expression have been previously explored^25^, we chose a method that closely resembles the native CRISPR–Cas system by flanking an array of gRNAs with ‘direct repeats’ (DR), which are the repetitive sequences required for the tandem gRNAs to be processed by RNase III in a tracrRNA-dependent manner^30–32^. Based on this approach, both gRNAs sg108 and LuxS1 were flanked with DR sequences and placed downstream of the *oxyS* promoter for peroxide-inducible control (**Fig. 4c**). We found elevated levels of AI-1 (**Supplementary Fig. 4d (i)**) and reduced levels of AI-2 (**Supplementary Fig. 4d (ii)**) in the peroxide-induced samples compared to the uninduced control, confirming CRISPR-mediated activation and inhibition on both QS signal producers. Surprisingly, we found that the cells that experienced more than one peroxide inductions displayed even lower AI-2 activity and higher AI-1 levels (**Supplementary Fig. 4d**). This result demonstrated dynamic control of CRISPR transcription regulation, hinting that the peroxide-mediated CRISPR activity could be ‘boosted’ by multiple applications of applied voltage over time to reach the desired target response. Correspondingly, dynamic multiplexed eCRISPR regulation is portrayed in **Fig. 4d**, in which we observed the highest AI-1 levels (2.2-fold increase) and lowest AI-2 activity (1.5-fold decrease) in the twice-induced electrode-bound bilingual cells. This again corroborated the fact that we can dynamically and electronically prompt the bilingual strain to transmit different types of signals and drive certain selected populations to elicit biological responses (e.g., bioluminescence). In sum, our results show a ‘bilingual’ *E. coli* strain capable of switching its QS signaling to reach different audiences based entirely on electronic cues.

## Communication with and automated control of biological activities

Through redox, “on-off” digital states of biological systems can be simply assigned and programmed with conventional electrochemical instrumentation and methods. Here, we demonstrate algorithm-facilitated, automated control of both enzymatic and eCRISPR activity in protein-based and cell-based subsystems. To achieve this, we designed custom electrochemical devices and developed models/algorithms to study, communicate with, and control the behavior of these biological subsystems.

First, we redesigned the above-noted 3D-printed optoelectrochemical device for encoding and decoding enzyme activity by regulating the activity of HRP through controlling the production of its substrate peroxide. We use electronics to ‘write’ chemical information that is ‘stored’ and then at a later time, electronically retrieved, enabling an electro-chemo-electro analog to the well-known random-access memory (RAM) of electronics. The custom electro-biochemical platform consists of a patterned ITO electrode that is separated into two interdigitated working electrodes and attached to a custom 3D-printed housing (**Fig. 5a-b**). Specifically, working electrode 1 (WE1) is tasked with generating peroxide (through ORR), hence ‘writing data’. Working electrode 2 (WE2), on which we electroassembled an HRP-conjugated gelatin hydrogel, is responsible for ‘reading’ the generated peroxide and ‘recording’ this information (in the form of electric current) (**Fig. 5b**). The recording function is enabled by measuring the current generated by the HRP-catalyzed enzymatic reaction with its substrate peroxide, and the subsequent redox cycling with mediator Fc (**Fig. 5c**)^17,20^. We found the endpoint current (recorded at 120 s) from WE2 increased each time a reducing charge was applied on WE1 (**Fig. 5d**), even with long periods (60 min) in between (**Fig. 5e, Supplementary Fig. 11**), confirming the platform’s ability to write, store, and retrieve data on enzyme activity.

**Figure 5.**
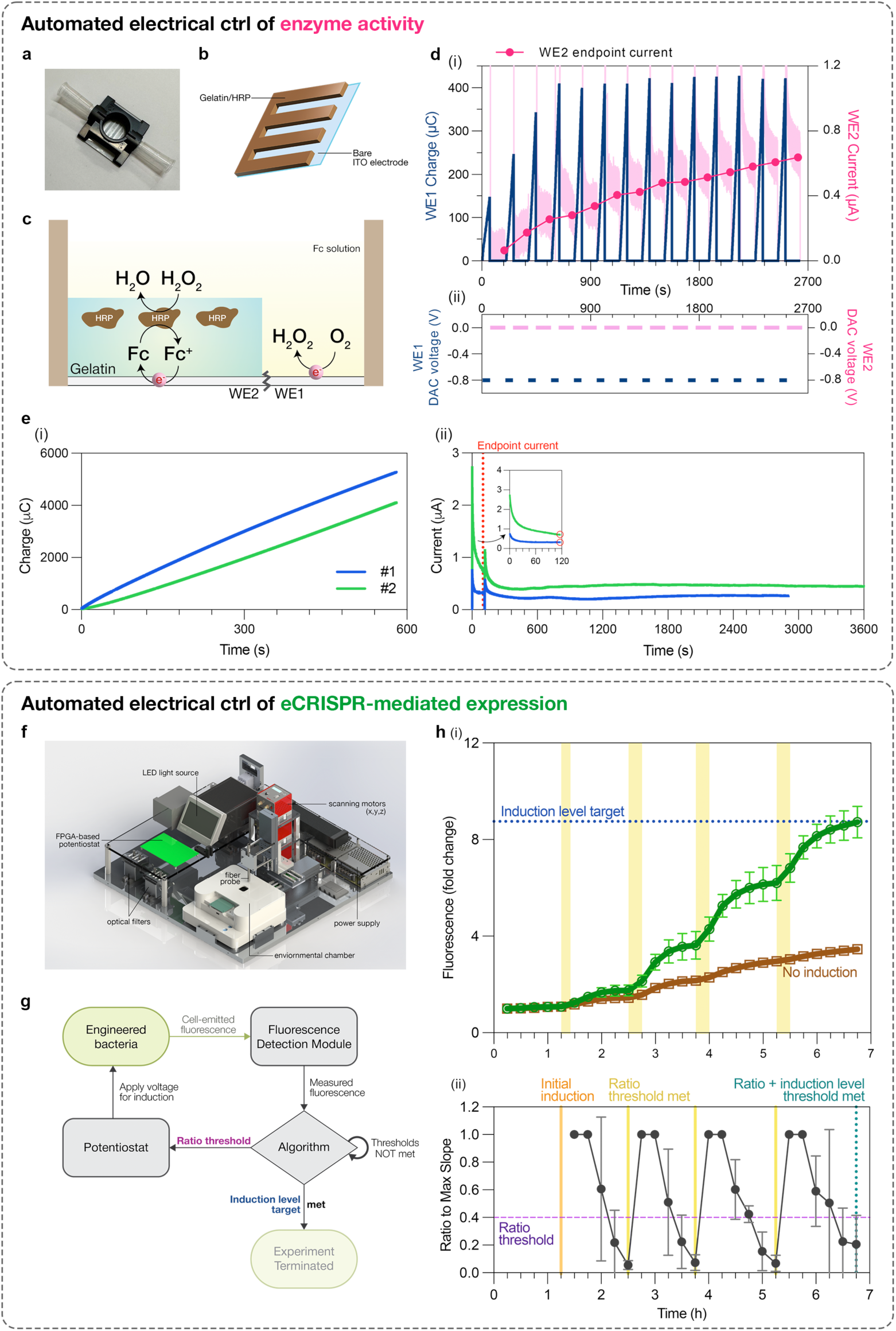
Automated dynamic control of biological activities. **(a)** Custom electrochemical platform with a patterned ITO glass slide attached to a 3D-printed housing. **(b)** An interdigitated electrode is generated through laser-cutting the ITO-coated glass into a zig-zag pattern to ensure the generated peroxide (on the bare electrode) is at the vicinity of the HRP/gelatin hydrogel for detection. **(c)** Working mechanism of the electro-biochemical ‘actuation checkpoint’ platform. A gelatin hydrogel containing horseradish peroxidase (HRP) is initially deposited on working electrode 2 (WE2). Working electrode 1 (WE1), consisting of just the bare ITO electrode, is tasked to generate peroxide when a reducing voltage is applied. The conjugated HRP in the gelatin gel on WE2 then catalyzes the generated peroxide, and the resulting electron cycles with the electrochemical mediator Fc that is present in the solution above. A current (from the cycling of Fc) can be read when an oxidizing voltage is applied (0 V vs Ag/AgCl). **(d)** Demonstration of data writing and storage. (i) Generated charge on WE1 (navy) and current obtained from WE2 (pink) were plotted over time. Pink filled circles represent the endpoint current recorded at 120 s. (ii) Voltages applied on WE1 and WE2 over time. **(e)** Demonstration of “long-term” data storage. (i) Total charge from applying −0.8 V on WE1 for 600 s. (ii) The following currents obtained for up to an hour on WE2 (inset: 0 – 120s). Red line and open circles indicate when the endpoint current was recorded (at 120 s). The blue dataset represents the first round (#1) of data writing (600 s) and recording (∼3000 s). The green dataset, representing the second round (#2) of data writing (600 s) and recording (3600 s), was conducted immediately after the first round. **(f)** Schematic illustration of the BioSpark system allowing electrochemical inducation and real-time fluorescence/electrochemical measurements. **(g)** Automation experiment workflow. Artificial biofilms containing engineered bacteria are assembled onto the ITO electrode of the optoelectrochemical device. The gene expression level, represented by the emitted fluorescence, is constantly monitored by ‘BioSpark’ and sent to the PC (diamond) for processing. Our custom algorithm (**Supplementary Fig. 7**) then determines the status of expression via calculating the rate of fluorescence increase (**Supplementary Methods**). When expression begins to tail, a user-specified threshold is met, and the potentiostat triggers an induction voltage for a specified charge. The experiment is automatically terminated when the fluorescence level is above a user-defined GFP threshold. **(h)** Automated dynamic control of eCRISPRa-regulated gene expression. (i) Fluorescence level within the artificial biofilm containing engineered eCRISPRa bacteria (NB101 harboring pSC-O108, pdCas9ω, and pMC-GFP). Fluorescence measurements were taken every 15 minutes (0.25 h). The experiment was terminated automatically when the fluorescence was above the specified threshold (blue dotted line). Brown open squares and line indicated the fluorescence level of a negative control to which no induction voltage was applied. Yellow zones indicate the duration of the applied voltage (−0.8 V). Note that the yellow zones also indicate the duration in which 0.4 ft^3^/h of oxygen was supplemented to both the experimental and negative control samples. A total charge of 2 mC was applied in each yellow zone. Open circles and open squares represent the mean, and error bars represent the standard deviation of individual replicates (*n* = 2 (biological replicates), and for each biological replicate three fluorescence measurements were performed and averaged). (ii) Ratio of slope, S, to S_max_ computed by our custom algorithm. Ratio threshold (purple dotted line) was set at 0.4. The orange line indicates when the algorithm applied the initial induction voltage. Yellow lines indicate when two consecutive ratios were below set threshold, thus meeting the ratio threshold and triggering the potentiostat to apply induction voltage. The teal dotted line indicates when both the ratio threshold and the induction level threshold were met, hence no voltage was applied, and the experiment was terminated. Filled circles represent the mean, and error bars represent the standard deviation of individual replicates (*n* = 2).

To realize automated control of *oxyRS*-based eCRISPR, we first studied the dynamic induction behavior of the *oxyRS* regulon when induced electrochemically. A simpler construct stripped of the CRISPR components but retaining its ability to respond to peroxide (encoded by *oxyS* promoter) was created. In suspension culture, we performed batch experiments and repeatedly induced with bolus additions of 50 or 100 mM peroxide. During both induction periods, we observed initial increases in GFP expression that were diminished after 45 minutes (**Supplementary Fig. 5a**), presumably due to transient depletion of peroxide^33^. We also show how cell growth was largely unaffected by repeated inductions (**Supplementary Fig. 5b**). These investigations exhibited that repeated induction could be sustained with minimal deleterious effect on cell proliferation and subsequent gene expression^18,34^.

Next, because optical (i.e., fluorescence) and electrochemical outputs either indirectly (optical) or directly (electrochemical) allow for real-time quantification that can be acted upon including by a process control scheme, they are both employed here as the primary signal modalities for bio-to-electronics communication. That is, to create a network for fully automated control, we constructed ‘BioSpark’, a complete bioelectronic system consisting of (i) a fluorescence detection module for gene expression measurements, (ii) a multichannel, FPGA-based potentiostat for sending and receiving electronic commands (and electrochemical detection), (iii) custom GUI program for system control and enabling local and wireless communication, and (iv) a custom 3D-printed environmental chamber for humidified, temperature-controlled, and oxygenated cell culture (**Fig. 5a** and **Supplementary Methods**). With BioSpark, multiple studies were carried out to examine the expression dynamics driven by on-device electroinduction. First, constitutive GFP-expressing *E. coli* helped us track its cell growth when assembled as artificial biofilms (**Supplementary Fig. 7a**). Using fluorescence levels as a surrogate for cell number, we observed a “lag-phase” of ∼1.25 hours immediately after electroassembly where no increase in fluorescence was detected. This lag period was consistently observed across many experiments, henceforth, we chose to initiate experiments, including electroinduction, with 1.25 h set as time zero. Next, NB101 cells harboring the peroxide-reporting plasmid (pOxy-sfGFP) were electrically induced (−0.8V) for different durations. Similar characteristics were observed to that of suspension cultures: (i) we saw an initial surge in electro-induced gene expression followed by a deceleration to a plateau (**Supplementary Fig. 7b**); (ii) this system retained its inducibility and repeatability, as shown by the increasing GFP levels when higher charge was placed (**Supplementary Fig. 7c**) and over multiple times (**Supplementary Fig. 8**). Based on these findings, we developed a phenomenologically-derived mathematical model of *oxyRS*-based electrogenetic expression (**Supplementary Methods**) so that a custom control algorithm (**Supplementary Methods**) could be implemented.

We then electroassembled the electrogenetic cells onto BioSpark’s electrodes and implemented a custom control algorithm, demonstrating for the first time, fully electronic feedback control of gene expression. **Fig. 5b** illustrates the automation workflow: gene expression, as represented by GFP levels, was measured and digitized via the fluorescence detection module to be relayed to the control algorithm. Based on many heuristic observations, we defined three model parameters that provide real time insight regarding the prevailing expression profile: a slope ratio that represented the change in the increase of fluorescence level, a threshold value for implementing simple proportional control, and a target value that served as an objective function (**Supplementary Methods)**. That is, we calculated in real time, the prevailing slope of the recorded fluorescence values. The maximum slope was updated continuously. We then computed the ratio between the prevailing and the maximum slope. If two consecutive ratios fell below the user-defined threshold, an additional potential was provided, and for consistency, each sequential pulse was of identical charge. Importantly, peroxide production is quantified by the prevailing current generated from the provided potential (**Supplementary Fig. 12-13**). These actions were repeated until later, when the target value was reached and the experiment was terminated (**Supplementary Fig. 7d**). With this simple control scheme in place, we then tested its efficacy. We assembled both peroxide reporters (**Supplementary Fig. 8**) and eCRISPRa cells (**Fig. 5c**) into ‘artificial biofilms’ and controlled their expression with the automated bioelectronic system. Simple visual examination of the profiles (**Figure 5h (i)**) shows how electroinduced cells rapidly increased their fluorescence immediately following the applied charge (within 15 minutes). Commensurate with both degraded signal molecule (peroxide^33^) and potentially reduced metabolic function^35^, the GFP fluorescence slowed and reached a plateau after an applied charge. In all cases (**Figure 5h (ii)**), our algorithm effectively identified each plateau region in the GFP expression profile and at each time signaled the potentiostat to apply a new charge (2 mC) enabling extra and consistent electroinduction^36^. Then, in **Figure 5h**, cells ultimately reached the target expression level (3.2×10^6^ AU) after four electroinductions (total 8 mC). Coincidently, in **Supplementary Fig. 12 and 13** we depict the current associated with electroinduction. As noted above, this electrochemical data dynamically reveals the rate of peroxide generation, which in turn, reflects the available dissolved oxygen at electrode surface. Given constant input of oxygen in a well-mixed system (**Supplementary Fig. 10**), this also indirectly indicates the respiratory activity of the cells immobilized in the hydrogel film. That the current was always above zero and monotonically decreased in time suggests both that the cells had oxygen and their continued growth in the films progressively inhibited its transfer to the electrode. In sum, these data demonstrate for the first time, closed-loop direct electronic control of gene expression on both direct peroxide-induced *oxyRS* expression as well as CRISPR-mediated *oxyRS* targeting^37^ enabled by the bioelectronic BioSpark system.

## Network integration for cascaded and supervised control

Biological systems, including living systems, are autonomously controlled locally as well as in cascaded networks via biological feedback (e.g., enzyme pathways with end product inhibition^38^, microbiome composition^39,40^, central and peripheral nervous systems^41^, the innate and adaptive immune systems^42,43^, including among seemingly disparate systems like the gut brain axis^44^). Owing to our ability to electronically survey, compute, and control biological systems, in **Fig. 6** we show how the above biological subsystems are used to create a network allowing multidirectional communication and cascaded, supervised feedback control of dynamic ‘living’ subsystems. The full network (as illustrated by the system diagram in **Supplementary Fig. 9**) is built upon the communication between (i) the BioSpark system for electrogenetic control of device-localized eCRISPR activity in cells, (ii) the bio-electrochemical platform for electric control of enzyme activity, and (iii) human users through the internet and text messaging. A simple algorithm was developed to control the electro-biochemical platform (**Fig. 6a**, denoted ‘remote’ system), which was then linked the ‘local’ electrogenetics system (BioSpark) for cascaded control of eCRISPR within the local BioSpark. As shown in **Fig. 6a**, local GFP expression levels of eCRISPR cells were monitored via BioSpark and these were relayed to the independently controlled bio-electrochemical platform for regulating HRP activity at a remote location. When the BioSpark algorithm detected a plateau in the GFP expression profile (again defined by the user-set ratio threshold (**Supplementary Methods**), a message was wirelessly conveyed to the remote bio-electrochemical platform over the internet. While there is no direct functional link between HRP activity and electrically-induced GFP expression in cells, the HRP serves here as a chemical/biological memory that is electronically addressed and in essence serves as an ‘actuation checkpoint’. Specifically, information pertaining to HRP’s enzymatic activity, as represented by the output current from the remote platform, is purposely designated to control the local BioSpark. In **Fig. 6b**, after the message from BioSpark was conveyed to the HRP electro-biochemical platform, its programmed procedure including the production of hydrogen peroxide (−0.8 V for 600 s on WE1) and retrieval of its level (0 V for 120 s on WE2) was then initiated. The resulting output current was then compared to another user-defined parameter (current threshold) that is used to feedback on and ultimately terminate eCRISPR activity at the local BioSpark. Initially at low peroxide level/HRP activity (**Fig. 6b**), this process served to cue the local BioSpark for additional electroinduction. Once the HRP activity had surpassed the threshold, the actuation checkpoint triggered a termination process for the BioSpark. This included both text messaging verification from human users on mobile phones and, once approved, the subsequent photobleaching of the electroinduced GFP (**Fig. 6b** and **Supplementary Figure 14**). In **Fig. 6b**, the photobleached reduction in fluorescence was observed. This demonstration, while there is no readily apparent biological basis for linkage (as is the case for feedback-controlled enzymatic pathways or innate/adaptive immunity), its example portends the future, where seemingly disparate biological systems can be studied, perhaps eventually linked and then even controlled.

**Figure 6.**
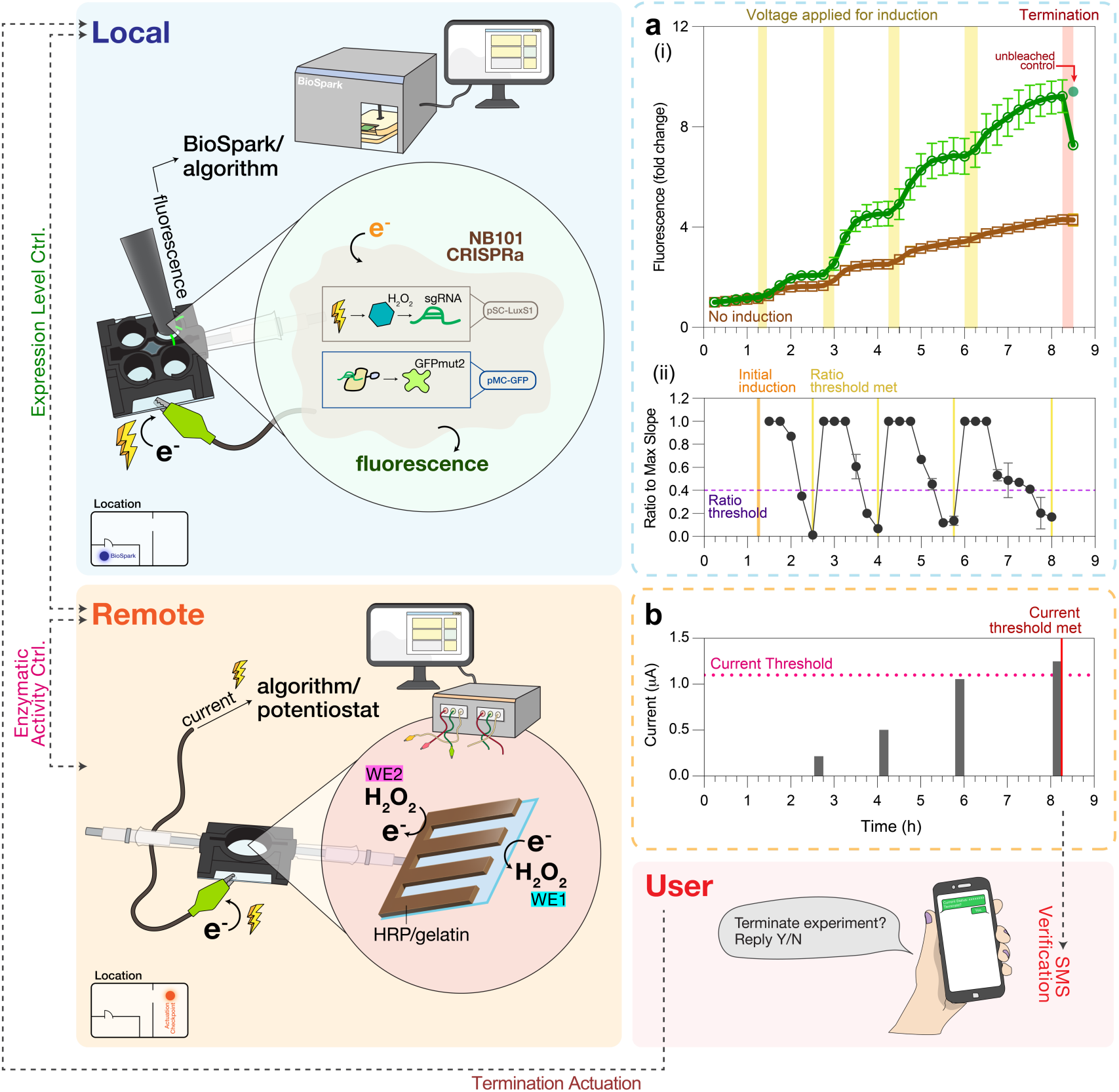
Network integration to enable remote feedback control of eCRISPR activity within the “Internet of Life”. **(a)** Automated feedback control of eCRISPRa-regulated gene expression (i) Fluorescence level of the artificial biofilm containing engineered eCRISPRa bacteria (NB101 harboring pSC-O108, pdCas9ω, and pMC-GFP, analogous to Fig. 5). Fluorescence measurements were taken every 15 minutes (0.25 h). Brown open squares and line indicate the mean in fluorescence level of the negative control to which no induction voltage was applied. Yellow zones indicate the durations when the induction voltage (−0.8 V) was applied to the experimental samples, and 0.4 ft^3^/h of oxygen was supplemented to both the experimental and negative control samples. A total charge of 2 mC was applied during each yellow zone. The red zone indicates when the selected individual sample was being photobleached. Open circles and squares represent the mean, and the green closed circle represent the unbleached control after termination. (ii) Ratio of slope, S, to S_max_ computed by our custom algorithm. Ratio threshold (purple dotted line) was set at 0.4. The orange line indicates when the algorithm applied the initial induction voltage. Yellow lines indicate when two consecutive ratios were below set threshold, thus meeting the ratio threshold and proceeded to trigger the remote ‘actuation checkpoint’. **(b)** Current threshold (pink dotted line) was set at 1.1 μΑ. The red line indicates when the current threshold was met, which prompted the PC at the remote location to send a text message to users.

## Conclusion/Discussion

In hopes of building an interconnected, feedback-controlled network of biological systems, we initially applied electro-biofabrication methods to assemble biological components in a manner that enables information exchange through redox. Enzymes and cells were placed in direct proximity to electrodes, with the latter forming “artificial biofilms”, a biomimetic structure. In this way, electro-biofabrication ensures efficient information flow at the bio-electronic interface and minimizes transport-limited heterogeneous responses of suspended biological systems^36,45^. Electroassembly opens avenues for immobilizing a multitude of living organisms, since our approach does not require extra protein or genetic engineering for assembly. This also relieves any extra metabolic burden associated with attachment and provides more effective use of resources for programmed functions^46^. Moreover, our assembly approaches are scalable and spatially programmable, inviting future opportunities for enzyme/cell grafting on complex formats like three-dimensional, miniaturized, arrayed electrodes; or on soft, flexible, bio-compatible materials, for wearable, ingestible or other portable systems^47,48^.

We then engineered *oxyRS*-based, peroxide-mediated eCRISPR that greatly expands the emerging repertoire of electrogenetics^7,11,49^. eCRISPR enables both upregulation and inhibition in a multiplexed manner for control of a variety of genetic targets. Here, we show that electronically programming two QS-related functions enables broad utility for manipulating microbial communities. Widespread in nature, QS signaling represents an ideal candidate for dispatching a highly-localized electrochemical cue to other *ex situ* biological populations with none or minimal genetic rewiring^28,50,51^, and achieving coordination within a natural or synthetic consortium^8,52^.

Next, automated electronic control of both enzymatic and genetic activities were demonstrated through device and algorithm development. While we have previously shown on-chip electronic modulation of chimeric proteins to control enzymatic activity during the generation of QS signals^18^, here we focused on the versatile redox-active enzyme, HRP. On top of electrochemical biosensors^53–55^, HRP is employed in many traditional, usually optical, biochemical assays^17,56–58^. Our current platform, with simple modification, could bring forth novel electrochemical procedures or even modular ‘bioproduction breadboards’ to interrogate or control biological processes^47,59,60^, especially considering that there is an extensive collection of redox-controlled enzymes. To realize electronic control of gene expression, responses from genetic activity were assessed using both light (GFP) and current (H_2_O_2_ generation) facilitating real time on-line control. We suggest that direct, electronic assessment and control augments the many transformative technologies associated with optogenetics^61–64^ and magnetogenetics^65^. By tapping into and controlling the native redox networks that are ubiquitous in biology, our approach offers a vast trove of targets and eventual applications. For example, eCRISPR provides a readily adaptable platform for simultaneous control of various genes and proteins via multiplexed transcriptional regulation. We further envision data analytics approaches such as machine learning, that when coupled with artificial intelligence, will greatly enhance phenomenological and first principles methodologies for optimized electrogenetic control.

Finally, in this work, we demonstrate wireless integration of bioelectronic systems forming a networked electrogenetic system. Our real-time, feedback control of eCRISPR activity was performed entirely electronically via redox signals and wireless internet connection/communication, suggesting convenient monitoring and feedback control of cells inside bioreactors for smart biomanufacturing^66^. Ultimately, all enzymatic and electrogenetics components in this study (that is, redox/peroxide signaling, CRISPR techniques, and QS communication) are either native to or can be easily ported to many biological systems; we foresee interconnected biohybrid networks that provide the foundation of a variety of intelligent systems and devices^67^. These include self-regulated, biomedical devices for *in situ* production and delivery of a therapeutic^68,69^, “smart” agricultural systems for rhizosphere manipulation^70^, or “sense-and-clean” strategies for battling environmental pollution^15,71^. Taken in sum, the concepts demonstrated here may serve as a blueprint for a more connected world in the future.

## Data Availability Statement

The datasets generated during and/or analyzed during the current study are available from the corresponding author on reasonable request. Source data are provided with this paper.

## Code Availability

The C# code for the gene expression algorithm in this study is available in the supplementary material or from the corresponding author upon request.

