## Supplementary Figures for "Redox-enabled Electronic Interrogation and Feedback Control of Hierarchical and Networked Biological Systems"

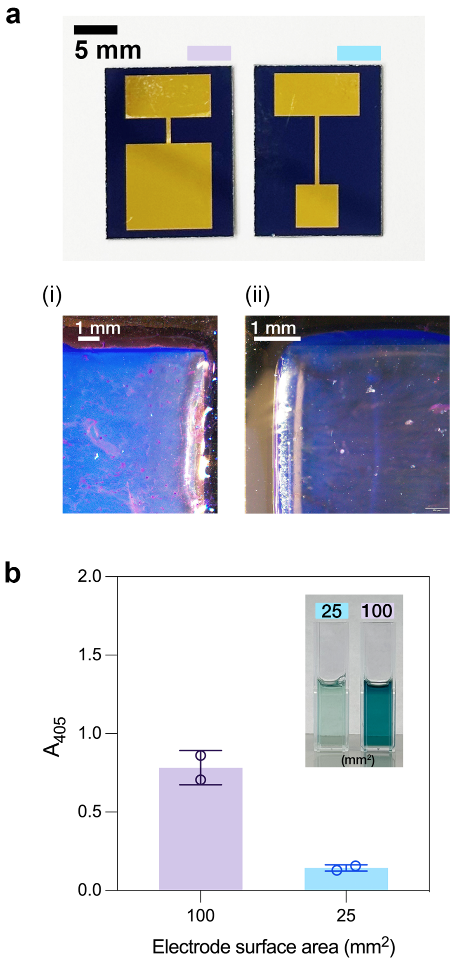


**Supplementary Figure 1. Spatially electrodeposited HRP/gelatin hydrogel. (a)** HRP/gelatin hydrogel electro-assembled on patterned gold electrodes with different surface area. (i) Brightview microscopy images, highlighting edges, of HRP/gelatin hydrogel stained with Coomassie Blue on a square 10 × 10 mm gold electrode. (ii) Brightview microscopy images of HRP/gelatin hydrogel stained with Coomassie Blue on a square 5 × 5 mm gold electrode. **(b)** Activity of electrode-bound HRP as determined via ABTS assay. Absorbance reading at 405 nm after 4 min of incubation with ABTS+$H_2O_2$ assay solution in room temperature. Bar height represent the mean, and error bars represent the standard deviation ($n = 2$). Individual replicates are plotted as open circles.

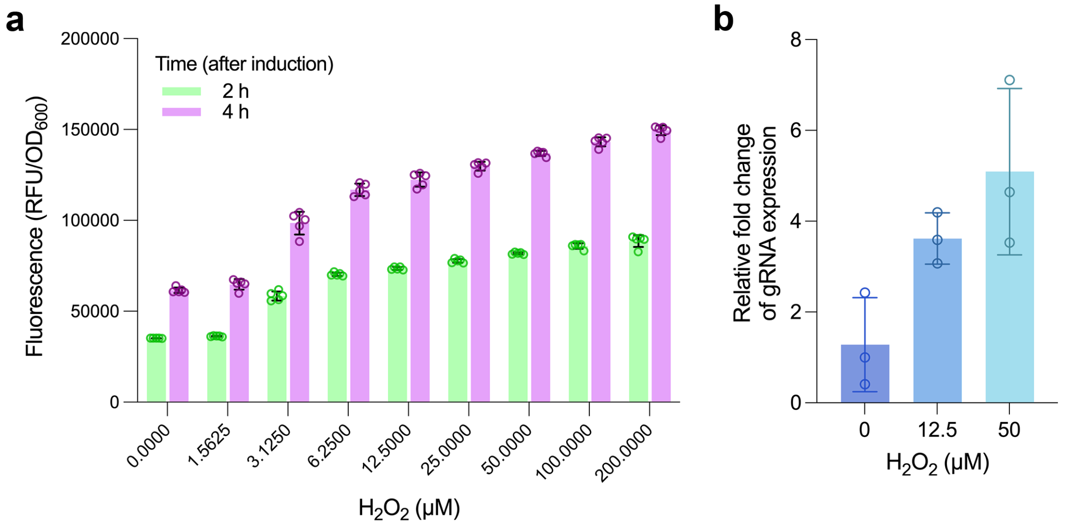


**Supplementary Figure 2. $H_2O_2$-inducible CRISPR activation of GFPmut2 (a)** Fluorescence of NB101 harboring pSC-O108, pdCas9ω, and pMC-GFP 2 hours (green) and 4 hours (purple) after $H_2O_2$ induction. Error bars represent the standard deviation ($n = 5$). Open circles represent individual replicates. **(b)** Relative fold change of gRNA (sg108) expression induced with various levels of $H_2O_2$. Error bars represent the standard deviation ($n = 3$). Open circles represent individual replicates. Bar height represents the mean for both figures.

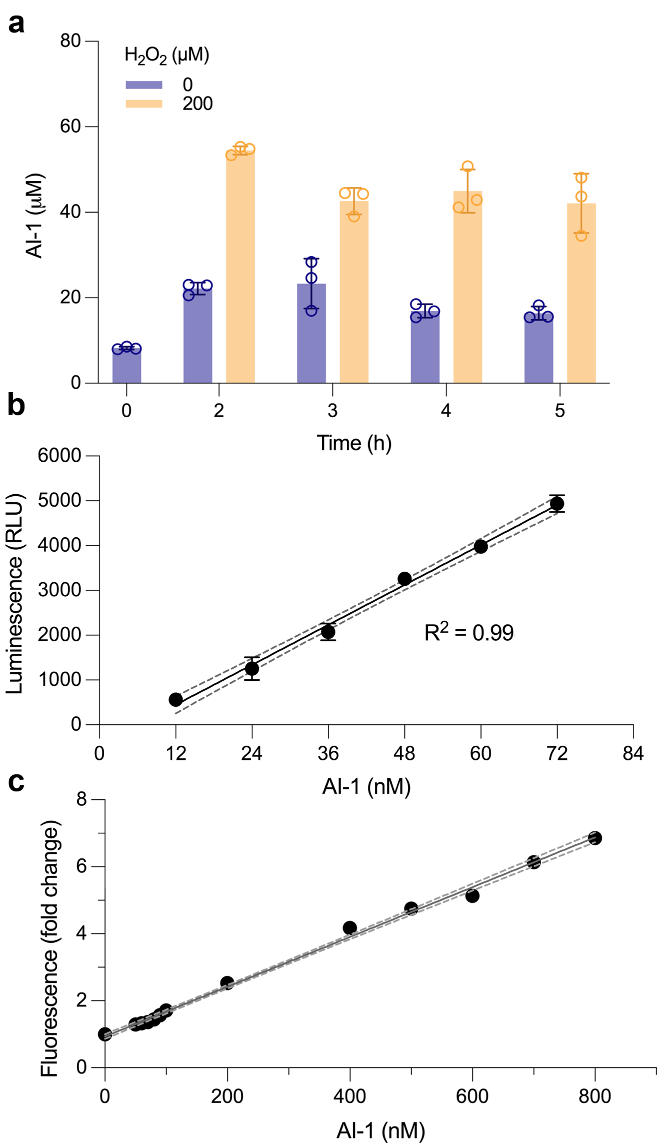


**Supplementary Figure 3. $H_2O_2$-inducible CRISPR activation of *lasI* (a)** AI-1 produced by NB101 harboring pSC-O108, pdCas9ω, and pMC-lasI-LAA after being induced by 0 µM (blue) and 200 µM (yellow) of $H_2O_2$. Bar height represents the mean value. Error bars represent the standard deviation ($n$ = 3). Open circles represent individual replicates. **(b)** AI-1 bioassay standard curve. Error bars represent the standard deviation ($n$ = 3). Calibration curve was generated via linear interpolation of experimental data, and the dashed lines represent the 95% confidence interval. **(c)** Normalized fluorescence response from AI-1 inducible strain (NEB10β + LasR_S129T-GFPmut3). Error bars represent the standard deviation ($n$ = 3).

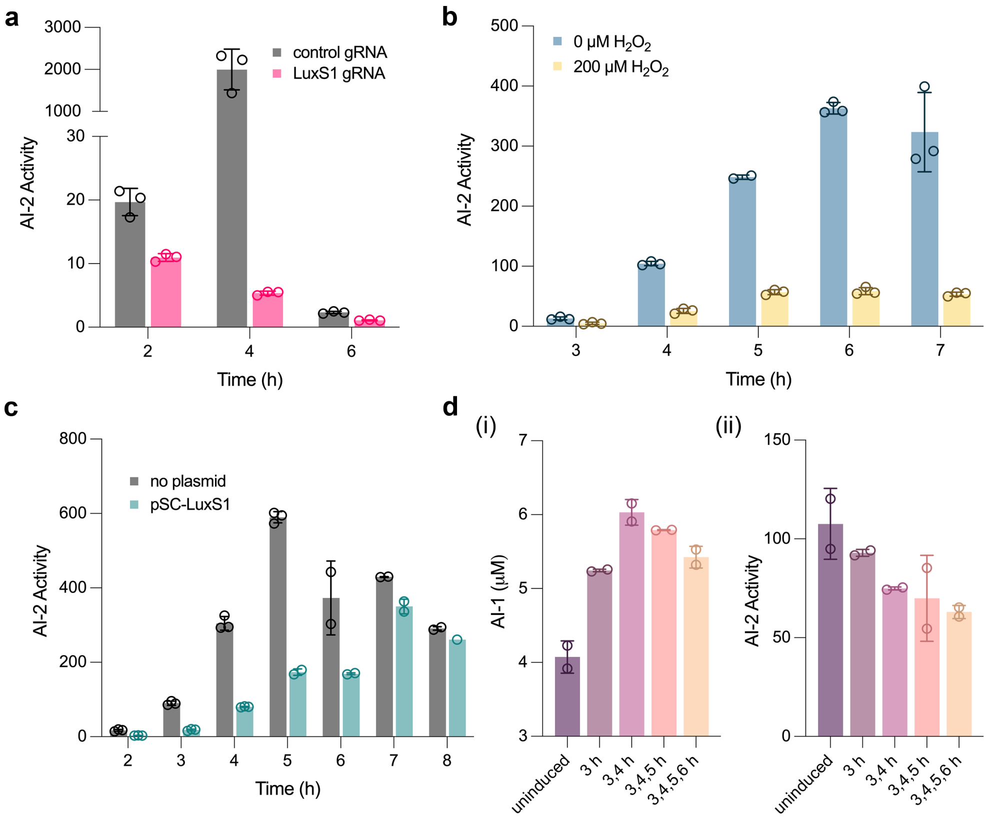


**Supplementary Figure 4. $H_2O_2$-inducible *luxS* CRISPRi and multiplexed control of QS signaling. (a)** Extracellular AI-2 activity of NB101 cells constitutively expressing either a control gRNA (grey) or *luxS*-specific gRNA LuxS1 (pink) at various time points after reinoculation. No glucose was added to LB media for inhibiting AI-2 uptake. **(b)** Extracellular AI-2 activity of NB101 cells carrying plasmids pSC-LuxS1 and pdCas9ω induced with 0 or 200 μM of peroxide at $OD_{600} = 0.4$. Conditioned media samples were collected at various time points after re-inoculation. A final concentration of 0.8% (w/v) glucose was added to LB media for inhibiting AI-2 uptake. **(c)** Extracellular AI-2 activity profile of NB101 cells harboring no plasmid (grey) or eCRISPRi components (pSC-LuxS1+pdCas9ω) when assembled as "artificial biofilms". **(d)** Measured AI-1 concentration (i) or AI-2 activity (ii) secreted from 'bilingual' cells 7 h post-reinoculation. Labels on the x-axis indicates when the cells received 200 μM $H_2O_2$ for inducing expression of both gRNAs. For all figures, bar height represents the mean and error bars represent the standard deviation (for **(a)** – **(c)** $n = 3$, for **(d)** $n = 2$). Open circle represents the individual replicates.

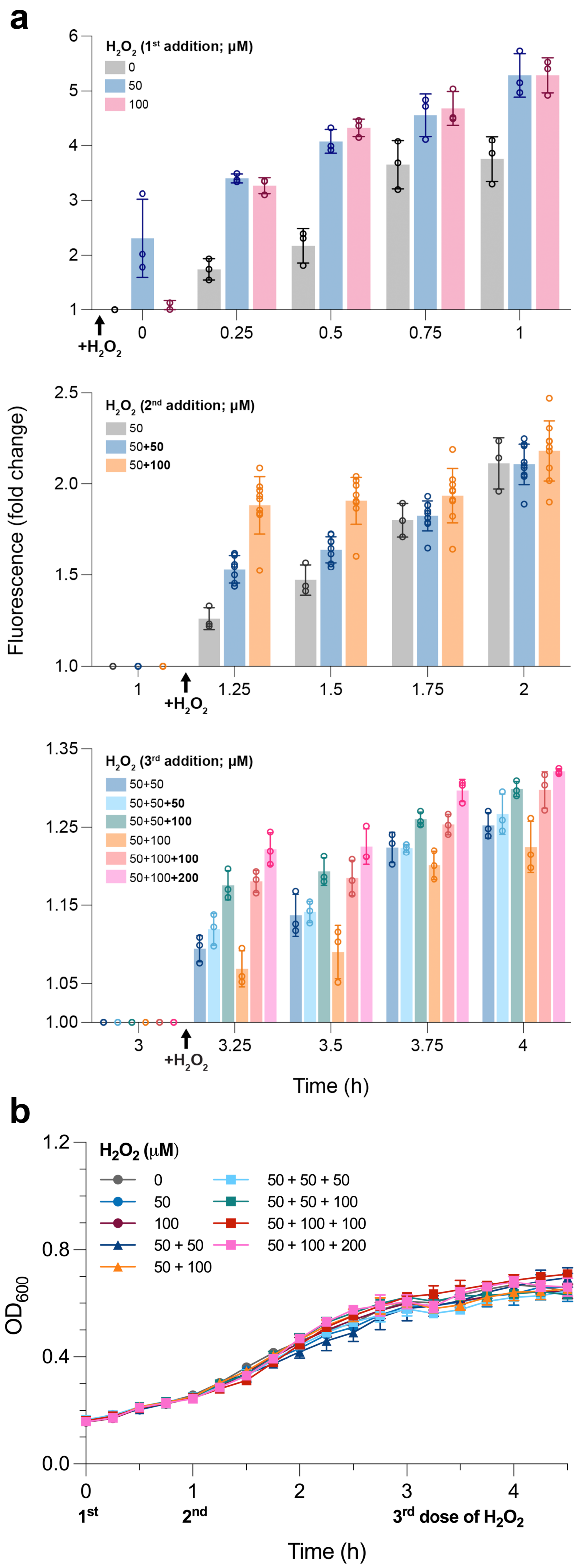


**Supplementary Figure 5. Peroxide-induced gene expression is dynamic and can be controlled with repeated inductions. (a)** Measured fluorescence from peroxide-reporter cells (NEB10β + pOxyRS-sfGFP-AAV). Cells were harvested at $OD_{600}$ = 0.4 via centrifugation and re-inoculated to $OD_{600}$ = 0.2 in 20% LB. Peroxide was added to each sample at indicated times (arrows). Fluorescence values from 0 - 1 h was normalized to the fluorescence of the uninduced control at time 0. Fluorescence values from 1 - 4 h was normalized to each sample's fluorescence at the beginning of each cycle (i.e., 1 or 3 h). Bar height represents the mean, and error bars represent the standard deviation ($n \geq 3$). Individual replicates were indicated by the open circles. **(b)** Growth curve of all experimental samples from repetitive inductions. Peroxide addition is indicated below its corresponding time. Filled circles (indicating one dose of peroxide), triangles (two doses), and squares (three doses) represent the mean. Error bars represent the standard deviation ($n = 3$).

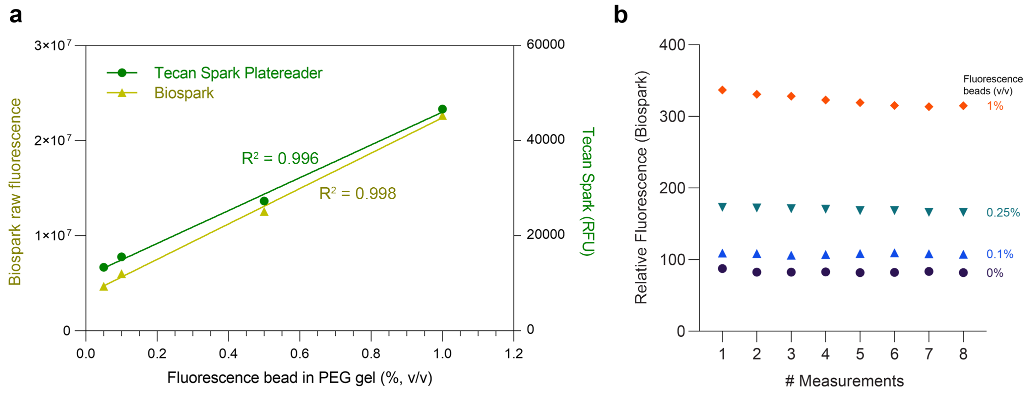


**Supplementary Figure 6. Fluorescence measurements of the BioSpark system. (a)** Green fluorescent particles were co-deposited with PEG-SH to various concentrations for simulating fluorescence emitted from engineered bacteria in the ‘artificial biofilm’. The resulted hydrogel was submerged in 150 µL of 20% LB, which is identical to the cell experiments. Fluorescence was read using BioSpark and the Tecan Spark® microplate reader. Linear responses for both were observed. **(b)** Relative measured fluorescence from fluorescent particles/PEG-SH hydrogel. The four samples were read in the sequence of 0% → 0.1% → 0.25% → 1% consecutively for 8 cycles.

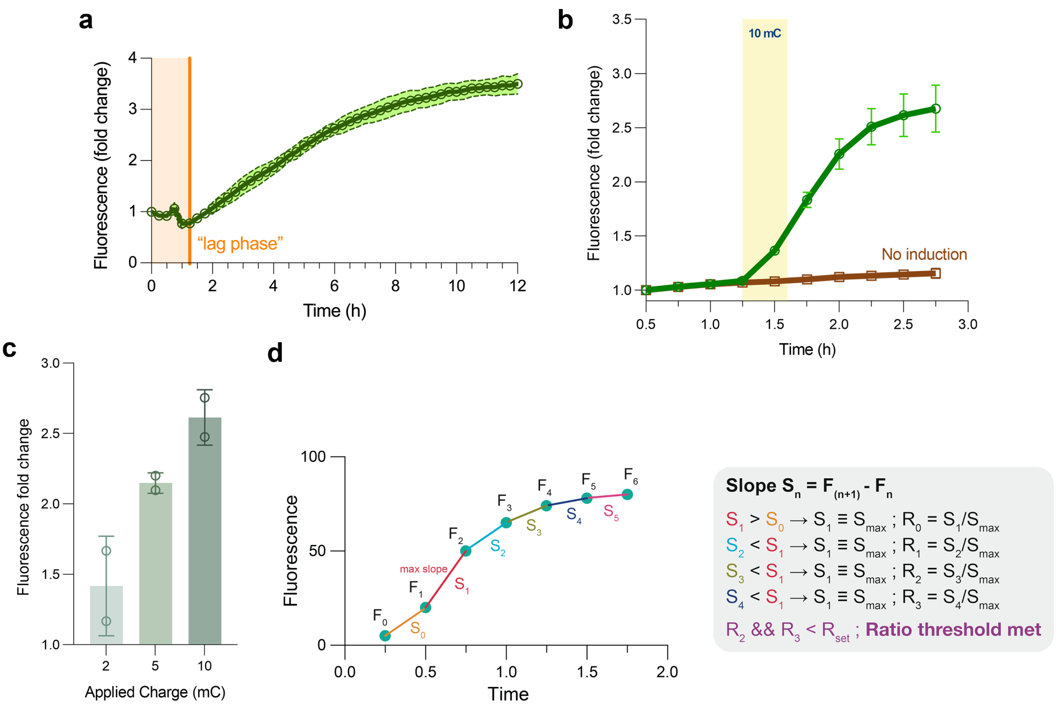


**Supplementary Figure 7. Development of a custom algorithm to achieve automated electrical control of gene expression. (a)** "Growth curve" of engineered *E. coli* entrapped in artificial biofilm from the measured fluorescence of DH5α-sfGFP cells co-deposited with PEG-SH. These cells constitutively express GFP in the absence of inducer; their expression is a surrogate for cell number. A "lag-phase" of approximately 1.25 hours was consistently observed during which there was no increase in fluorescence, suggesting that there was no growth during this period. Open circles represent the mean fluorescence, and the dashed curves represent the error from individual replicates ($n$ = 4). **(b)** Representative gene expression dynamic for in-film peroxide reporters (NEB10β + pOxy-sfGFP). The yellow zone (vertical band) indicates period during which voltage was applied for electroinduction. The uninduced negative control is also shown (in brown). Note that there are four wells in the device operated in parallel (hence two biological replicates for both experimental sample and negative control). Each well is interrogated with a moving optical fluorescence probe. Fluorescence from each well is measured three times and the average is reported. The elapsed time of the entire process (to measure fluorescence for all four wells) is typically less than two minutes. Open circles and squares (indicated) represent the mean, and error bars represent the standard deviation of individual biological replicates ($n$ = 2). **(c)** Fold change in fluorescence before and after electroinduction. Bar height represents the mean and error bars represent the standard deviation ($n$ = 2). Individual replicates are indicated by the open circles. **(d)** Basis data pairing for phenomenological algorithm underpinning gene expression assessment (see **Supplementary Methods**). Since the fluorescence measurements were taken at fixed time intervals, we defined the slope as the difference between two neighboring fluorescence measurements. The algorithm stores and updates the value of the maximum slope. It also computes the ratio between the current slope and the maximum slope ($S_{max}$). If two consecutive ratios fall below the user-set limit (expressed as a ratio), the algorithm considers the threshold met and initiates electro-induction via the potentiostat. The $S_{max}$ will, in turn, return to 0 and a new cycle will begin.

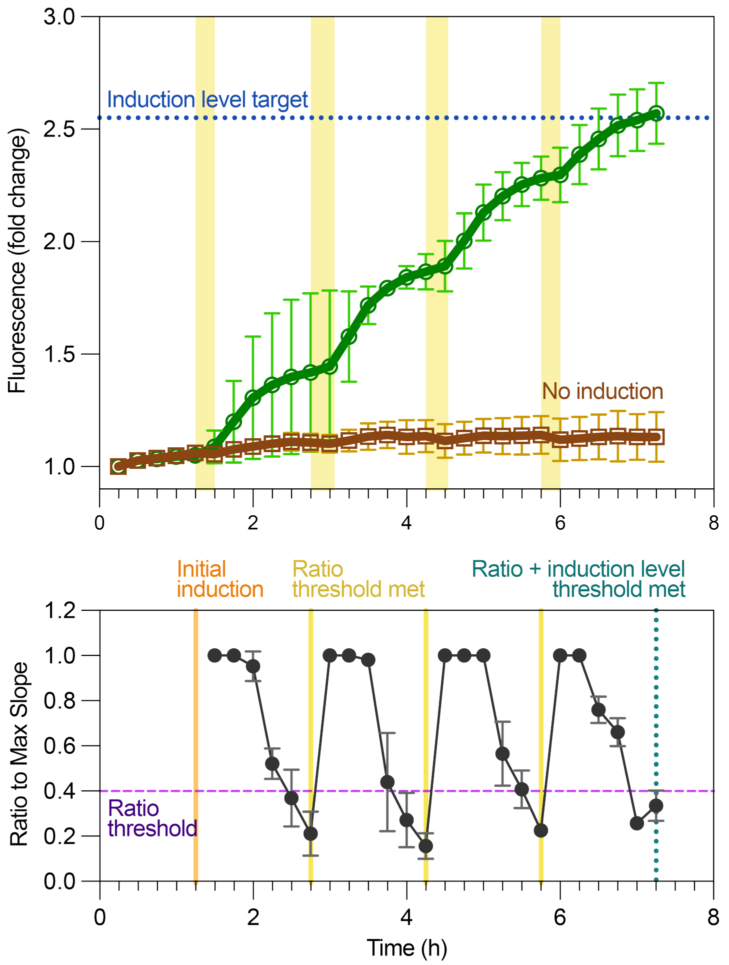


**Supplementary Figure 8. Automated dynamic control of electro-induced gene expression.** Top: Fluorescence level of the artificial biofilm containing peroxide reporters (NB101+pOxy-sfGFP). The experiment was automatically terminated when the fluorescence exceeded the induction level target (blue dotted line). The brown dataset indicates the fluorescence level of the negative control to which no induction voltage was applied. Yellow zones indicate the duration over which the induction voltage (-0.8 V) was applied. Note, to provide sufficient peroxide and metabolic activity, each experimental and negative control samples was supplemented with 0.4 $ft^3$/h of oxygen (flow rate determined over many runs). A total charge of 2 mC was applied for each induction (width of yellow zone indicates the amount of time to reach 2mC). Open circles and squares represent the individual biological replicates.

Bottom: Ratio of slope, S, to $S_{max}$ computed by our custom algorithm. Ratio threshold (purple dashed line) was set at 0.4. The orange line indicates when the algorithm applied the initial induction voltage (1.25 h). Yellow lines indicate that two consecutive ratios were below the set point threshold, thus meeting the ratio threshold and triggering the potentiostat to apply induction voltage. The teal dotted line indicates when both the ratio threshold and the induction level threshold were met, hence no voltage was applied, and the experiment ended. For all figures, the error bars represent the standard deviation of individual replicates (*n* = 2).

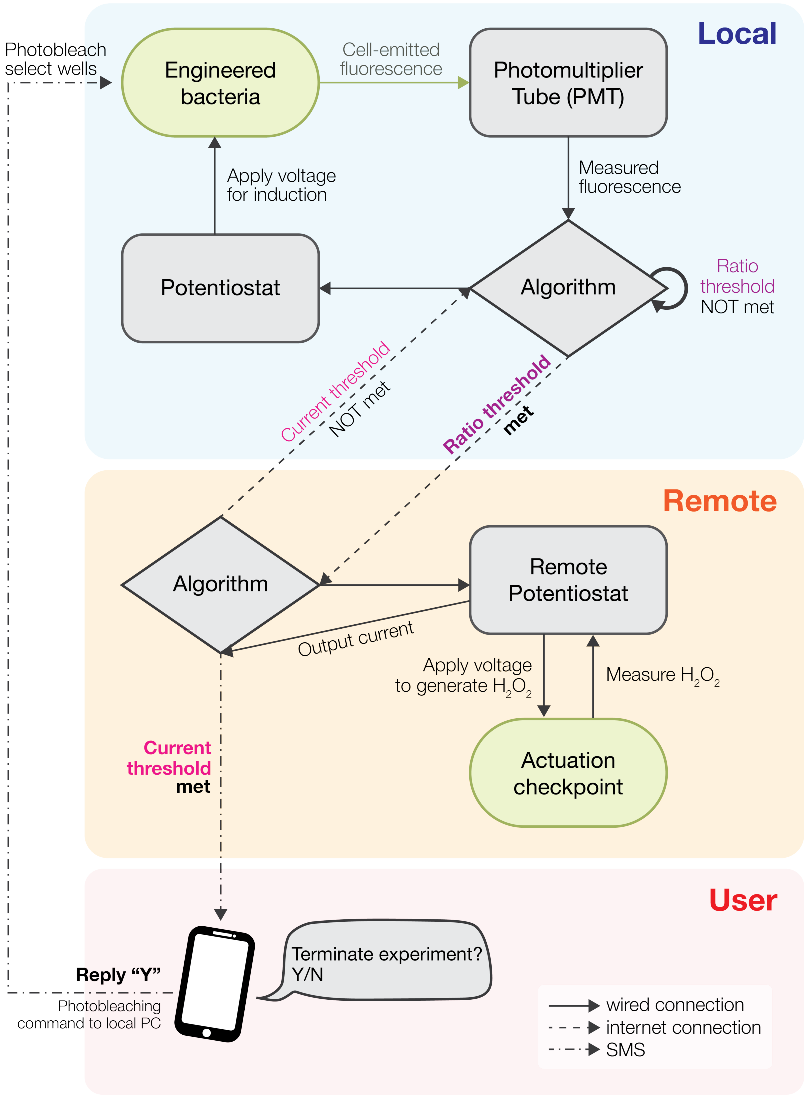


**Supplementary Figure 9. System diagram of the integrated network representing the “Internet of Life”.** *Local:* Emitted cell fluorescence is measured three times (over 2 minutes) by the fluorescence module in the BioSpark system every 15 minutes and fed to the custom algorithm for fluorescence/expression level tracking. The local algorithm (see **Supplementary Figure 7**) (i) stores and updates the $S_{max}$, (ii) computes the slope ratio, and (iii) compares the current ratio to the user-defined ratio threshold. When two consecutive slope ratios are less than or equal to the threshold value, the local algorithm considers the ratio threshold met and sends a message to the PC situated at the remote location wirelessly through the internet.

*Remote*: When the bio-electrochemical platform (connected to a PC) (denoted “actuation checkpoint”) located at the remote location receives the message from the local BioSpark system, the remote algorithm then commands the remote potentiostat to run a pre-set program. -0.8 V will first be applied on WE1 for peroxide generation, followed by 0 V on WE2 for peroxide detection. The remote algorithm then compares the value of the output current to that of the user-defined current threshold: if the output current does not exceed the threshold, a message is returned (via internet) to the local BioSpark system to administer electroinduction immediately after the upcoming fluorescence measurement (for a user-defined duration or charge). The elapsed time for this occurrence was typically 15 minutes. Otherwise, the remote algorithm sends a SMS verification message to alert the human users and seek permission to terminate the experiment.

*User*: Human users can respond to a query by sending text messages on a mobile phone (by replying “Y “or “N”) to indicate whether the experiment should be terminated or not. If replied with “Y”, BioSpark initiates the photobleaching program as a demonstration to destroy the expression product.

### Supplementary Notes (for Figures 2-5)

#### *Figure 2*

In **Figure 2f**, we examined the difference in current obtained during electroinduction (-0.8 V) with or without exogenous oxygen supplement. Surface-assembled *E. coli* (NB101 + pOxy-sfGFP, $OD_{600}$ = 6 $ml^{-1}$), submerged in 20% LB, was acclimatized in the custom environmental chamber at 34°C for 1.5 hours prior to electroinduction. The blue curve shows the output current without exogenous oxygen supplement (~0.5 µA, indicating minimal $H_2O_2$ generation), while the purple curve shows the current during which we supplied 0.4 $ft^3$/h (~0.01 $m^3$/h) of oxygen (starting from 0 s) through built-in tubing in the connector of the 4-well biohybrid electronic device. In the case of supplemented oxygen, it took about 2 minutes, but the measured current was observed to increase steadily. This profile is likely due to the active metabolism of the *E. coli* cells entrapped in the artificial biofilm that depleted the dissolved oxygen in the culture media. After two minutes, increased current shows that oxygen was present at the electrode surface, suggesting sufficient oxygen for respiratory function. Considering that the current for the controls without cells (both gel and no-gel) was always higher than the gel with cells, this indicates that the cells did indeed metabolize provided oxygen. Thus, we designed the 3D-printed biohybrid electronic device to include built-in tubing for exogenous oxygen supply. Schematic illustrations of the entry ports are shown in **Supplementary Figure 10**. We believe that both oxygen diffusion and the gentle stirring (convection) caused by the airflow facilitated oxygen transport, and together, sufficient oxygen could be delivered to the electrode surface allowing both cell respiration and electroinduction.

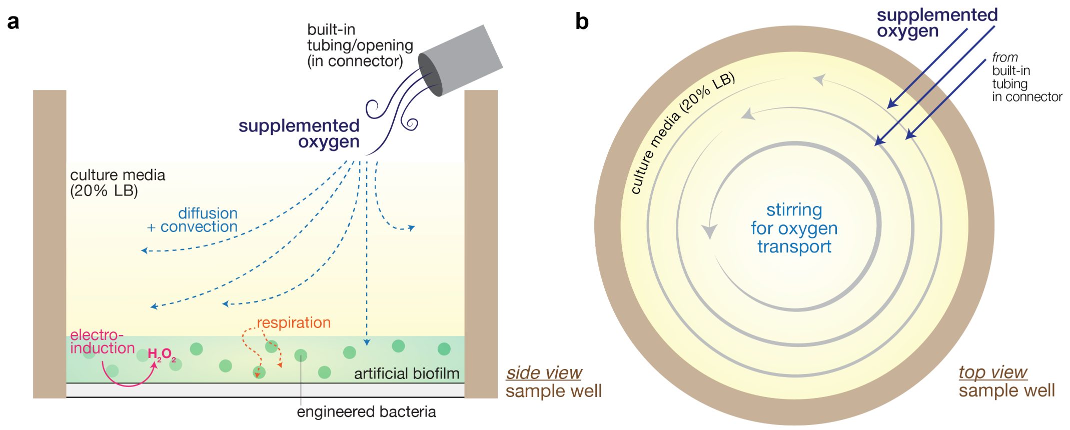


**Supplementary Figure 10. Schematic of the mechanism for exogenous oxygen supply. (a)** Side view of an individual sample well in the biohybrid electronic device. **(b)** Top view of an individual sample well in the biohybrid electronic device.

***Figure 3***

In **Figures 3d-f**, we reported the total applied charge for each individual experiment. We believe the charge variations from the experiments mainly resulted from the difference in electrode surface area, the slight potential deviation from the reference electrodes, and the content of the culture media.

***Figure 4***

In **Figure 4d**, we too reported the total applied charge for the electroinduced samples. For the 0 h and 0&3 h samples, we applied -0.8 V for 30 minutes (resulting in a charge of 2.71 mC) to both immediately after deposition. After 3 hours of incubation at 37°C, we administered an additional 30 minutes of -0.8 V electroinduction to the 0&3 h sample (resulting in a charge of 0.43 mC). We believe the charge (or accordingly, current) difference reflects the decrease of available oxygen in the media, since the cells were actively respiring, and no oxygen was supplemented to the samples in this experiment.

***Figure 5***

In **Figure 5e**, we showed that the information "written" by WE1 (i.e., generated peroxide) can be "stored" in our electro-biochemical device and "retrieved" by WE2 at a later time. For this, we performed data writing (on WE1) and recording (on WE2) for a prolonged period (**Supplementary Fig. 11**). First, -0.8 V was applied to WE1 for 600 s to generate peroxide. 0 V was then applied to WE2 for peroxide detection, and we observed minimal decrease in WE2 current throughout the entire recording period (from 600 s to 3600 s for the #1 round). Next, this process (both data writing and recording) was repeated immediately (denoted the #2 round) and we saw an increase in WE2 current (throughout the entire 3600 s recording period), indicating that the increased peroxide level (from the #2 round of data writing) was stored and detected.

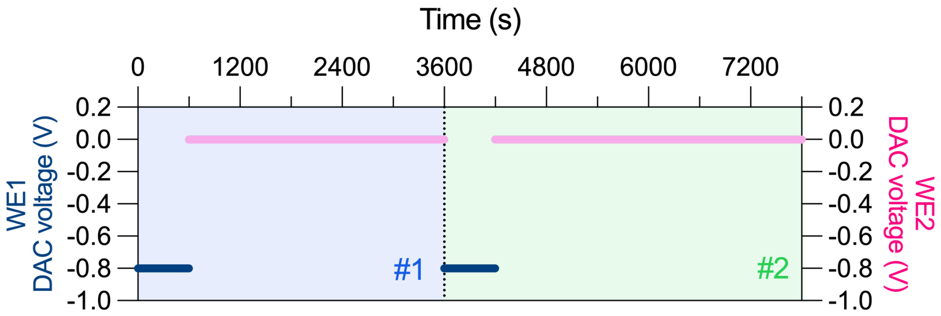


**Supplementary Figure 11**. Voltages applied on WE1 and WE2 of the electro-biochemical device (from **Fig. 5e**).

In **Figure 5h (i)**, the thickness of the yellow zones demonstrates the time over which electroinduction was administered (i.e., peroxide was generated). During this period, we also supplied 0.4 $ft^3/h$ of oxygen to both the experimental samples and the negative controls. Since the total charge in each yellow zone was kept constant (2 mC), the increased time (from ~12 min to ~20 min) of voltage (-0.8 V) application to reach this charge also reflects the decreasing oxygen level at the electrode surface. As shown in **Supplementary Figure 12**, the generated current within each electroinduction cycle is dynamic and increases along the electroinduction period due to the supplied oxygen. We note also that the total current gradually decreased as the cycles progressed. We believe this reflects a decreasing oxygen level at the electrode commensurate with increasing cell number (**Supplementary Figure 7a**) present in the artificial biofilm. On the other hand, the transient increases in fluorescence from the uninduced negative control may be due to added in oxygen availability from the oxygen line since oxygen was only supplemented during electroinduction periods.

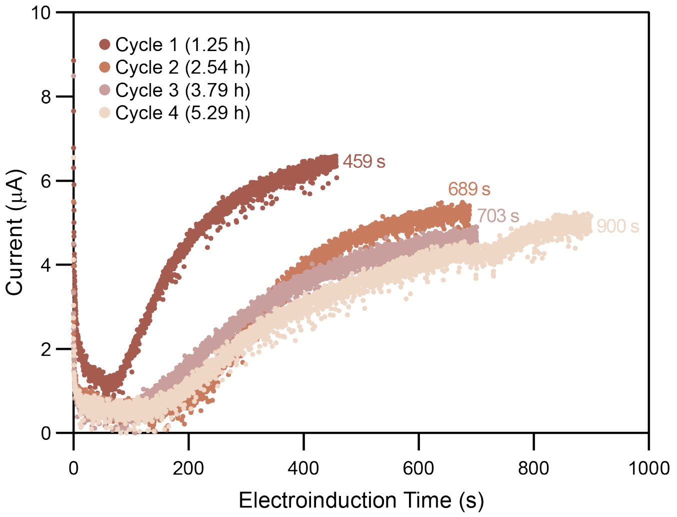


**Supplementary Figure 12.** Generated current during electroinduction (from **Figure 5h** (i)).

#### *Figure 6*

In **Figure 6a (i)**, like **Figure 5h (i)**, the thickness of the yellow zones demonstrates the time over which electroinduction was administered (i.e., peroxide was generated). These datasets are from two independent experiments but are very similar in the result. Like in **Figure 5h**, during this period, 0.4 $ft^3/h$ of oxygen was supplied to both the experimental samples and the negative controls. We observed similar electroinduction and expression behavior here: (i) the increase in electroinduction duration along the time suggests the decrease in oxygen level at the electrode surface; and (ii) the transient increases in fluorescence from the negative control might also be due to changes in oxygen availability since oxygen was only supplemented during electroinduction. The generated current during electroinduction is shown in **Supplementary Figure 13**. Here again, the observed duration of the 2 mC charge increased in time as the culture progressed to the higher fluorescence levels and cell densities.

To terminate the experiment, we used the fluorescence measurement module in BioSpark to photobleach the GFP. Schematic illustrations of the two different functions of the fluorescence measurement module are shown in **Supplementary Figure 14**. For fluorescence measurements, a pulsed (0.07 ms) excitation light (at 469 ± 17.5 nm) was directed to the samples for 100 times and concurrently the optical probe also collected the emission light (at 525 ± 19.5 nm). For photobleaching, a continuous excitation light (469 ± 17.5 nm) was directed to the samples (for 15 min) instead.

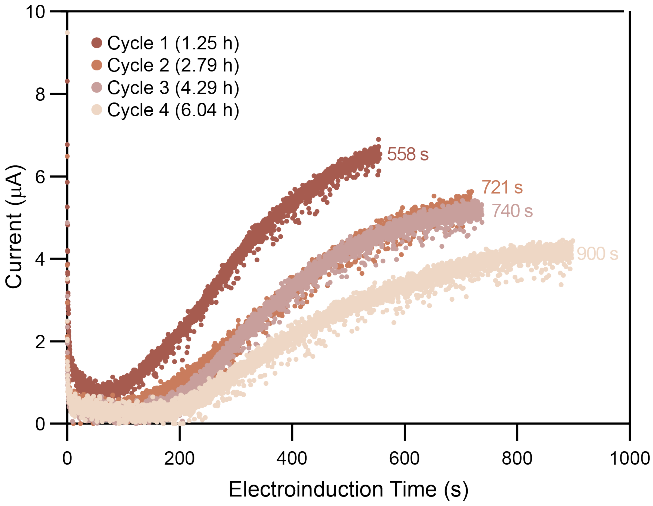


**Supplementary Figure 13.** Generated current during electroinduction (from **Figure 6a** (i)).

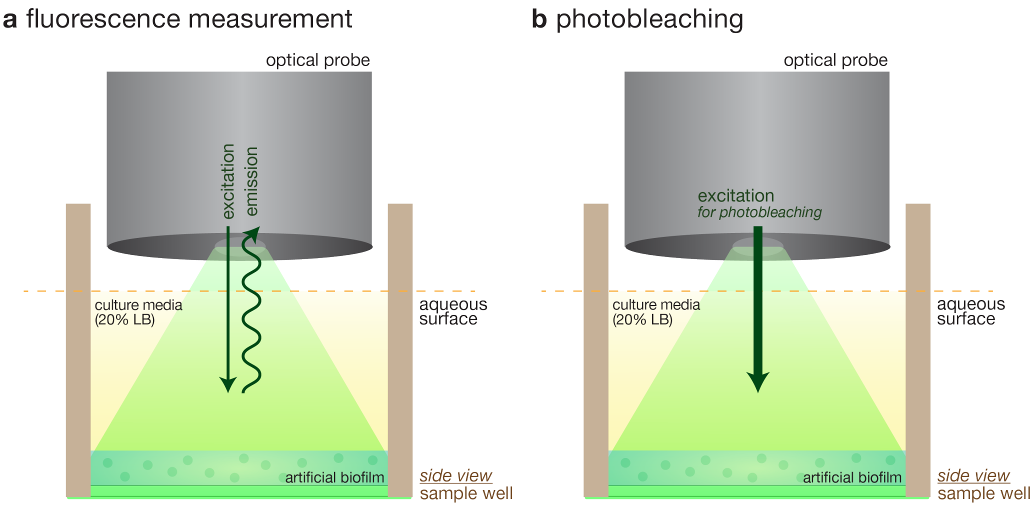


**Supplementary Figure 14. Schematic of the fluorescence measurement module in BioSpark. (a)** Fluorescence measurements **(b)** photobleaching
