## Supplementary Materials for "Redox-enabled Electronic Interrogation and Feedback Control of Hierarchical and Networked Biological Systems"

**Supplementary Materials (excluding Supplementary Figures) for** Redox-enabled Electronic Interrogation and Feedback Control of Hierarchical and Networked Biological Systems

**Authors:**

Sally Wang[123*], Chen-Yu Chen[123*], John R. Rzasa[2], Chen-Yu Tsao[23], Jinyang Li[23], Eric VanArsdale[123], Eunkyoung Kim[23], Fauziah Rahma Zakaria[123], Gregory F. Payne[23], William E. Bentley[123]

1 Fischell Department of Bioengineering, University of Maryland, College Park, Maryland, USA

2 Fischell Institute of Biomedical Devices, University of Maryland, College Park, Maryland, USA

3 Institute of Bioscience and Biotechnology Research (IBBR), University of Maryland, Rockville, Maryland, USA

**Table of Contents:**

Methods……………………………………………………………………………………….2

Supplementary Methods………………………………………………………………………5

Supplementary Tables..………………………………………………………………………9

References………….....……………………………………………………………………..12

### Methods

##### *Chemicals*

Potassium chloride (KCl), $H_2O_2$ (30%), phosphate buffered saline (PBS), potassium phosphate monobasic ($KH_2PO_4$), potassium phosphate dibasic ($K_2HPO_4$), potassium hexachloroiridate(III) ($K_3IrCl_6$, Ir), and gelatin from porcine skin (gel strength ≈175 g Bloom, Type A, G2625) were purchased from Millipore-Sigma. All antibiotics (ampicillin, kanamycin, and chloramphenicol) were purchased from Millipore-Sigma. Lysogeny broth (LB) and agarose were from Fisher Scientific. Fluorescent beads and Pierce® horseradish peroxidase (HRP) were purchased from Thermo Fisher Scientific. 4-arm PEG-SH (MW 5000) were from JenKem USA. 1,1'-Ferrocenedimenthanol (Fc) was from Santa Cruz Biology. AI-1 (N-3-oxo-dodecanoyl-L-Homoserine lactone) was from Cayman Chemicals. Fc and AI-1 were initially prepared as 1 M and 5 mM DMSO stocks.

##### *Culture media and conditions*

Unless otherwise indicated, cells were grown overnight in LB at 37 °C, 250 r.p.m. shaking, inoculated at $OD_{600}$ = 0.1 in LB media the following day, and grown until the indicated cell density (optical density at 600 nm, $OD_{600}$). Optical density was measured using an UV-Vis Spectrophotometer (Beckman Coulter).

##### *Plasmid construction*

All bacterial strains and constructs used in this study were listed in **Supplementary Table 1**. All enzymes, competent cells and reagents were from New England Biolabs and used according to provided protocols. Q5 polymerase and primers in **Supplementary Table 3** (gene sequences in **Supplementary Table 2**) were used for PCR reactions. DpnI digestion, polynucleotide kinase phosphorylation, T4 ligations, Gibson assembly and *E. coli* chemical transformation were performed using New England Biolabs product protocols. DNA clean-up (Zymo Research), gel extraction (Zymo Research) and plasmid preparation kits (Qiagen) were performed using provided protocols. Synthetic gene fragment containing the multiplex crRNAs was purchased from Thermo Fisher Scientific. Synthetic gene fragment containing the 108 spacer and gRNA scaffold was purchased from Integrated DNA Technologies (IDT). Synthetic gene fragment containing tracrRNA was purchased from Integrated DNA Technologies (IDT).

##### *Electrodeposition of E. coli and PEG-SH*

*E. coli* grown to mid-log phase were harvested via centrifugation at 3000 rcf for 10 minutes, then resuspended in 1× PBS to 2× the desired OD. To prepare the 2× PEG-SH solution, 100 mg/mL of PEG-SH were dissolved in phosphate buffer (PB, 0.1 M pH = 7.4) containing 10 mM Fc, as described previously by Li *et al*[1]. Prior to electrodeposition, the two solutions were mixed at a 1:1 ratio. We then performed chronoamperometry, poised at 0.8 V for 30 s (or stated otherwise), to initiate oxidation of the thiol group for crosslinking. The endpoint charge was recorded for each run. Excess cell/PEG-SH solution was then removed, and the generated film was carefully washed with PBS to remove any unbound cell/PEG-SH.

##### *Fabrication of the biohybrid electronic device*

The custom-designed device was designed based on a standard 3-electrode system. The resin-based housing was fabricated with a Mars 3 Pro 3D printer with the standard black resin from ELEGOO (Guangdong, China) and was then attached to a 25 mm × 25 mm ITO-coated glass substrate (Sigma) as the working electrode using via photo-curing with the same resin for 3D printing. To cast the agarose salt bridge, the solution containing 0.1% agarose in 1 M KCl was first heated and then pipetted into the central well. After the agarose solidified, 1 M KCl was added to submerge the salt bridge. The Ag/AgCl reference electrode (Pine Research) can be connected to the central well containing the salt bridge via the side opening. A separate custom connector was also fabricated using the 3D printer and resin described above. This connector comprised a platinum wire as the counter electrode and four air nozzles designed for evenly distributing oxygen to the independent wells for later hydrogen peroxide production. With the device fully assembled, both the inserted reference electrode and the counter electrode would be immersed in the 1 M KCl solution present in the central salt bridge well. Together with ITO working electrodes, this system formed a complete circuit for the electrochemical operation. (**Fig. 1d**).

***Quantification of film thickness***

Overnight culture of DH5α-sfGFP was harvested via centrifugation (3000 rcf, 15 min) and resuspended in PBS to $OD_{600} = 2\ mL^{-1}$. 2× PEG-SH solution was prepared as previously stated and was mixed 1:1 with the *E. coli* solution. 100 µL of cell/PEG-SH mixture were loaded into the wells of the biohybrid electronic device and electrodeposition (0.8 V) was performed for the indicated duration. After decanting the excess solution and thorough washing, a ZEISS LSM700 confocal microscope was used to obtain Z-stack images of the generated film. Using a 5 µm Z-stack depth, we defined the film thickness based on the distance between the electrode and the boundary layer (the last Z-stack slice with visible *E. coli*).

***Peroxide Generation and Quantification***

Electrochemical peroxide generation was performed in the biohybrid electronic device with its setup as described above. 150 µL of 20% LB mixed with 80% PBS (henceforth, 20% LB) was added into each well (surface area = 38.5 $mm^2$). The working electrode solution was undisturbed (or, where indicated, stirred via blowing $O_2$ into the wells). The electrodes were connected to a potentiostat (either 700-series CH Instruments or a custom FPGA-based potentiostat). Chronoamperometry, poised at −0.8 V for the indicated duration, was performed to generate hydrogen peroxide. The endpoint charge was recorded for each run.
To quantify the generated peroxide, we used the Pierce® quantitative peroxide assay kit (aqueous) (Thermo Scientific) according to the manufacturer's instructions. Briefly, the working reagent was prepared by mixing one volume of Reagent A with 100 volumes of Reagent B, with at least 200 µl prepared for each sample to be assayed. Ten volumes of the working reagent were added to one volume of sample (typically 200 µl working reagent to 20 µl sample) in a well of a clear-bottomed 96-well plate. The reaction was mixed and incubated for 15–20 min, after which a Spark® microplate reader (Tecan) was used to measure the absorbance at 595 nm. Sample peroxide concentration was calculated by comparison with a standard curve (dilutions of 30% (*w/w*) peroxide) performed on the same day.

***General and coculture electroinduction set-up***

Electroinduction experiments were performed in the custom biohybrid electronic device. Deposition of the cell/PEG-SH film was performed as described in previous sections. After thorough washing to remove excess cell/PEG-SH solution, 150 µL of 20% LB was added to each well as the culture media. Peroxide was generated via chronoamperometry as described in sections above, with voltage application for a specified duration (e.g., 1800 s). The biohybrid device was then moved to an incubator (30 or 37°C, as indicated) for incubation. For all automated experiments, the biohybrid device remained in a custom-made environmental chamber inside the Biospark system with temperature (34°C) and humidity (≥ 80%) control. 0.4 $ft^3/h$ (scfh) of oxygen was supplied to all samples (including the negative control) during electroinduction for media perturbation and oxygen supply to generate sufficient peroxide.
Samples (i.e., the media immersing the film) were removed at indicated time intervals and sterile filtered for downstream bioassays. For coculture experiments, AI-1 responsive strain (NEB10β + pLasR_S129T-GFPmut3) were grown to mid-log phase and reinoculated to an $OD_{600}$ of 0.025 in 20% LB. Culture was then pipetted into the wells of the biohybrid device in which the 'artificial

biofilm' (containing CRISPRa *lasI* cells) was previously deposited. Electroinduction was subsequently performed as described above.

***AI-1 quantification***

AI-1 quantification was performed by bioluminescence assay. AI-1 reporter cells JLD271 pAL105[2] were grown overnight in LB at 37 °C and 250 r.p.m. shaking with the appropriate antibiotics. The following day, AI-1 solutions (0 – 84 nM) for the standard calibration curve were prepared in 20% LB. The reporter cells were diluted 500-fold in LB with the appropriate antibiotics. For every experimental replicate, 90 µl of diluted reporter cells and 10 µl of the standard AI-1 solutions were added into the wells of a white-bottom 96-well plate (Corning). Experimental conditioned media samples were prepared similarly after sterile filtering and diluting between two- and thousand-fold to maintain a linear assay range. Microplate with reporter cells and conditioned media samples were incubated at 30 °C and 250 r.p.m. shaking in a Tecan® microplate reader, and its luminescence was measured by the plate reader every 30 minutes for 3-5 hours. The AHL concentration of each sample was calculated using the standard curve.

***AI-2 activity assay***

Relative AI-2 levels were determined using the *V. harveyi* reporter BB170 bioluminescence assay[3] with slight modifications. AI-2 reporter cells BB170 were grown overnight in AB media at 30°C and 250 r.p.m shaking with appropriate antibiotics. The following day, overnight BB170 culture was diluted 5000-fold in AB media. For every experimental replicate, 180 µl of diluted BB170 culture and 20 µl of conditioned media were added into the wells of a white-bottom 96-well plate (Corning). The microplate was then incubated at 30 °C and 250 r.p.m. shaking in a Tecan® microplate reader, and its luminescence was monitored by the plate reader every 30 minutes after 3-hour incubation. AI-2 activity was calculated by dividing the RLU produced by the reporter after addition of conditioned media by the RLU of the reporter when growth medium alone was added.

***Electrodeposition of HRP/gelatin hydrogel and electrochemical detection of $H_2O_2$***

We followed the protocol for HRP/gelatin deposition and electrochemical detection of peroxide as described by Li *et al.* with slight modifications[1]. Deposition of HRP/gelatin was performed on a custom patterned electrode generated through laser-cutting the ITO-coated glass slide into two separate interweaving working electrode. This zig-zag pattern was chosen to ensure the generated peroxide be at the vicinity of the deposited HRP/gelatin hydrogel for detection. A custom 3D-printed housing with one central well exposing the working electrodes and two side openings for the insertion of Ag/AgCl reference electrodes was then attached to the patterned ITO electrode (**Fig. 5a-b**). A custom connector attached with two Pt wires provided the 3-electrode system with counter electrodes. Upon deposition, a solution containing HRP (1 mg/mL in 0.1 M PB, pH = 6), gelatin (25 mg/mL), and $Ir^{3+}$ (5 mM) was pipetted into the central well of the device followed by applying an oxidative voltage (+1.2 V) for 2 minutes on working electrode 2 (WE2). After removing the excess HRP/gelatin mixture, a solution of 0.25 mM Fc in 0.1 M PB (pH = 7.4) was added into the well submerging both working electrodes. Electrochemical peroxide generation was performed on working electrode 1 (WE1) by applying -0.8 V for the desired duration as indicated. To detect the peroxide generated from WE1, a constant voltage (0 V vs Ag/AgCl) was imposed on WE2 for 120 s, and the endpoint current output was recorded (**Fig. 5d-e**).

### Supplementary Methods

#### A. Experimental Methods

***Electrode chip fabrication***

Gold patterned electrodes were purchased from Platypus Technologies and Pine Research. Steps for fabricating the custom patterned gold electrode on silicon wafers by Platypus Technologies were as follows: First, metal deposition was performed on standard 4-inch silicon wafers using a Denton thermal evaporator (Denton Vacuum), with metal deposition rates of 2–3 Å $s^{-1}$. Specifically, a 50 nm chromium adhesion layer was evaporated, followed by 100 nm gold. Next, photolithography utilized direct writing of photoresist via a DWL66fs laser writer (Heidelberg Instruments), guided by a laser exposure map designed in AutoCAD (Autodesk). Photoresist spin-coating and development steps were performed using an EVG120 automated resist processing system (EV Group). The patterned wafer was post-processed by etching, photoresist stripping and cutting individual electrodes with a DAD dicing saw (DISCO). The patterned gold electrode purchased from Pine research was made by screen-printing gold on a ceramic base, with a 2 $mm^2$ diameter gold working electrode, a printed Ag/AgCl reference electrode, and a printed gold reference electrode.

***Visualization of the electrodeposited hydrogels***

Gold patterned electrodes were submerged in a solution containing HRP (0.1 mg/mL in 0.1 M PB, pH = 6), gelatin (25 mg/mL), and $Ir^{3+}$ (5 mM) for electro-deposition (+1.1 V, 1 min). The resulting hydrogel was then stained with Coomassie Blue solution (0.1% Coomassie® R-250 in 50% ethanol and 10% acetic acid) followed by brief destaining (50% methanol and 10% acetic acid). Brightview images of the hydrogel was taken by the MVX10 upright fluorescence microscope (Olympus). Overnight BL21 culture was adjusted to OD ~ 12 in 0.85% NaCl solution and stained with SYTO-9 dye (final concentration 8.35 μM) (Thermo Fisher Scientific) for 15 minutes in the dark. The stained cultures were then mixed 1:1 with 50 mg/mL PEG-SH + 10 mM Fc solution for deposition (+0.7 V, 2 min) on gold patterned electrodes. Both fluorescence and brightview images were taken by the MVX10 upright fluorescence microscope (Olympus).

***Colorimetric HRP enzymatic activity assay***

Gelatin/HRP hydrogel was deposited on two gold patterned electrodes with different surface areas (25 or 100 $mm^2$) as described in the section above. The resulting hydrogel was then washed in 0.1 M PB (pH = 6) for 10 minutes before the enzymatic assay. A final concentration of 0.8 mM 2,2'-azino-bis(3-ethylbenzothiazoline-6-sulfonic acid (ABTS) and 0.003% (0.88 mM) $H_2O_2$ as assay solution mixture was prepared from an ABTS stock solution (16 mM) and 30% $H_2O_2$. The washed hydrogel (on gold patterned electrode) was incubated in 2.2 mL of the assay solution mixture for 4 minutes at room temperature. The hydrogel was then removed and 1 mL of the assay solution mixture was transferred to a 1-cm cuvette for absorbance reading at 405 nm using an UV-Vis spectrophotometer (Beckman Coulter).

***Cloning of peroxide-inducible gRNA expression plasmids***

Plasmid pSC-O108 (**Supplementary Table 1**) for peroxide-inducible expression of sgRNA sg108 was made via PCR amplification and Gibson assembly. First, pSC-108gRNA[4] was removed of *soxR*, *soxRS* promoter region, and sgRNA sg108 by restriction digestion with ClaI and BamHI. Gene fragment containing *oxyR* and *oxyS* promoter (**Supplementary Table 2**) was amplified from plasmid pOxy-LacZlaa[5] with primers SW01 and SW02 (**Supplementary Table 3**). After DpnI treatment, fragments were ligated by Gibson Assembly. Next, gene fragment containing spacer 108 and gRNA scaffold was then PCR amplified with primers SW03 and SW04 and inserted into the linearized intermediate product via primers SW05 and SW06 to generate the construct pSC-O108.

Plasmid pSC-LuxS1 for peroxide-inducible expression of sgRNA LuxS1 was made via site-directed mutagenesis. Spacer 108 (**Supplementary Table 2**) in plasmid pSC-O108 was swapped with LuxS1 (**Supplementary Table 2**) using primers SW07 and SW08.

Plasmid pSC-sg108+LuxS1 for peroxide-inducible expression of crRNAs spacer 108 and LuxS1 was made vis PCR linearization, restriction digestion, and T4 ligation. pSC-O108 was initially PCR linearized using primers SW09 and SW10 to remove the spacer 108 and the gRNA scaffold, as well as adding restriction sites BamHI and XhoI. Gene fragment containing crRNAs spacer 108 and LuxS1 (**Supplementary Table 2**) was inserted into linearized pSC-O108 backbone via restriction digestion and T4 ligation. After this, the gene fragment containing the tracrRNA (**Supplementary Table 2**) was inserted into the product generated from the previous step via restriction digestion and T4 ligation to generate the final construct.

***Cloning of pLuxS1***

Plasmid pLuxS1 for constitutive expression of sgRNA LuxS1 was constructed via site-directed mutagenesis using primers SW11 and SW12 to swap the spacer 108 in plasmid pS108gRNA[4] to LuxS1.

***Cloning of pMC-lasI-LAA***

Plasmid pMC-lasI-LAA was constructed via PCR amplification and Gibson assembly. pMC-GFP[4] was linearized and removed of GFPmut2 by primers SW13 and SW14. Primers SW15 and SW16 were used to amplify and add the LAA ssRA tag to *lasI*. *lasI* with the added LAA tag was then inserted to the linearized backbone via Gibson assembly.

***Cloning of pOxy-sfGFP-AAV***

Plasmid pOxy-sfGFP-AAV was constructed via site-directed mutagenesis using primers SW17 and SW18 to add the AAV ssRA tag to pOxy-sfGFP[6].

***Fluorescence measurements***

Unless stated otherwise, we used a Spark® plate reader (Tecan) to measure GFP fluorescence with excitation/emission wavelengths of 488/520 nm. For dynamic gene expression control experiments, GFP fluorescence was measured using our custom-built Biospark platform, as described in later sections.

***RT-qPCR***

NB101 harboring the plasmids that allow CRISPR activation of GFP (pSC-O108 + pdCas9ω + pMC-GFP) were grown at 37°C and 250 r.p.m in LB overnight. The following day, overnight cultures were diluted to $OD_{600}$ = 0.1 and grown at 37°C and 250 r.p.m until reaching mid-log phase. Peroxide stock solution (10 mM) were then spiked into the cultures to induce the expression of sgRNA. After 2 hours of incubation, samples were collected via centrifugation. Total and microRNA were extracted using a miRNeasy Kit (Qiagen) and quantified using a Nanodrop (Thermo Scientific). RT-qPCR was then carried out using primers SW19/SW20, the Power SYBR™ Green RNA-to-CT™ 1-Step Kit (Applied Biosystems), and a Quantstudio 7 Flex Real-time PCR system (Applied Biosystems).

#### **B.** (Local) **Biospark System for gene expression control**

***Custom-built bioelectronic system setup***

Hardware of the Biospark system includes a custom two-channel arbitrary waveform potentiostat with simultaneous sampling of up to two PMT sensors, a motorized fiber-coupled fluorescence reader, and a custom environmental chamber. This system is controlled by a custom Windows GUI program.

**(1)** *Fluorescence measurement and photobleaching module*

The fluorescence module consists of a 470nm excitation light source and driver (Thorlabs #M470F4 and #DC2200) that sent light to a fiber-coupled filter mount (Thorlabs #FOFMS) via a 1000μm diameter core multimode fiber (Thorlabs #FT1000EMT-CUSTOM). An OD6 fluorescence filter (Edmund Optics #67-027) was used to set the excitation band. This light was then directed onto the test target via a fiber coupled reflection probe (Thorlabs #RP22). The emission from the target was collected by the other fibers in this bundle and directed to a second fiber coupled filter mount (Thorlabs #FOFMS) which contained two OD6 emission filters (Edmund Optics #67-030). Two filters were used to improve out-of-band blocking with only a modest reduction in the emission signal. The filtered light in turn passed through another fiber (Thorlabs #FT1000EMT-CUSTOM) into a fluorescence enhanced PMT (Thorlabs #PMT2101) where it was converted to an electrical signal that travelled to the analog-to-digital converter (ADC) on the FPGA potentiostat board. A 3-axis motion platform consisting of three stepper motor kits (Thorlabs #KMTS50E) connected together with a base plate (Thorlabs #MTS50A-Z8), XY plate (Thorlabs #MTS50B-Z8) and right-angle plate (Thorlabs #MTS50C-Z8) was used to allow x, y, and z-axis movement of the probe. To administer photobleaching, we used the fiber-coupled reflection probe to direct emission light to the sample well for a desired duration (for 15 min in the experiment showcased by **Figure 6**). A schematic depicting the setup during fluorescence detection and photobleaching is shown in **Supplementary Figure 10**.

**(2)** *Custom FPGA-potentiostat*

We designed and fabricated a custom FPGA-based two channel potentiostat PCB controller. The core component is a Spartan-7 module (Opal Kelly #XEM7305) and breakout board (Opal Kelly #BRK7305). Power was supplied via an ultralow-noise linear power supply (Acopian #DB5-50). While many components are used on the custom potentiostat board, the three primary components of interest include the (a) femtoampere input bias current op-amps (Analog Devices #ADA4530), (b) the 16-bit multichannel Digital-to-Analog converter (DAC) (Analog Devices #AD5765), and the 18-bit multichannel ADC (Texas Instruments #ADS8698).

**(3)** *Power source, connection to PC, and custom control software*

General power for the Biospark system is provided from four power supplies: a 5VDC (CUI # VGS-75C-5), a 12VDC (CUI # VGS-75W-12), a 15VDC (CUI #VGS-75C-15), and a 48VDC (CUI #VGS-75W-48). The system is connected to a PC via USB (Latorice #B07GLP8SD). A custom Windows GUI program written in C# is used to control the system. Functions included adjusting metrics related to the (i) various potentiostat modes (e.g., chronoamperometry mode, cyclic voltammetry mode, manual electroinduction mode, algorithm-controlled electroinduction mode), (ii) potentiostat settings (e.g., voltage, pulse period, slew rate, maximum total charge, etc...), (iii) fluorescence reading (e.g., pulse width, pulse count, measurement interval), and (iv) automated algorithm control (e.g., algorithm/manual control mode, ratio thresholds, expression level target, SMS settings, etc...). Both electrochemical and fluorescence data are saved in a custom binary format but can be exported into an Excel spreadsheet (.xlsx). On the main screen of the program, four real-time diagrams displayed the fluorescence values, potentiostat currents and charges, applied counter voltages, and the CV curve or fluorescence slope ratios. The entire Biospark system is housed in an opaque enclosure to ensure no stray light enters the test chamber.

***Custom algorithm for "smart" control of gene expression***

We established a simple model-based algorithm (generic model control)[7] to monitor gene expression for the surface-assembled recombinant *E. coli* producing either transcription or translation products as represented by the emitted fluorescence (F). The process was described by a dynamic model:

$$F = g(x, u, t) \tag{1}$$

where **x** is the state vector, **u** is a vector of potentiostat inputs, and *t* is time. Process input *u(t)* can be described by Equation 2:

$$u = h(a, t) \tag{2}$$

where **a** is the current resulted from electroinduction, and *t* is time. An algorithm was then developed from this model to track the output fluorescence and control the system (**Supplementary Figure 7**):

Since the fluorescence were taken at a fixed time interval (15 min/0.25 h), we defined the slope (S; RFU×4×$h^{-1}$) as the difference between two neighboring fluorescence measurements as described by Equation 3.

$$S_n = F_{(n+1)} - F_n \tag{3}$$

The algorithm stores and updates the value of the maximum slope ($S_{max}$) to serve as a reference point for the progression of the fluorescence level. The change in fluorescence levels was determined by the ratio (R) between the current slope and $S_{max}$, as described by Equation 4.

$$R_{(n-1)} = S_n \div S_{max} \tag{4}$$

If two consecutive ratios fall below the user-defined ratio value (set as 0.4 in both **Figure 5** and **Supplementary Figure 8**), the algorithm considers the ratio threshold is met and will initiate electroinduction by sending a command to the potentiostat. $S_{max}$ will return to 0 and a new cycle will start subsequently. Variables such as the ratio threshold, duration of voltage application, or the total charge applied can be set in the GUI program.

#### **C.** (Remote) **Enzymatic Activity Control Checkpoint**

##### ***Hardware, software, and connection to PC***

An individual custom two-channel arbitrary waveform potentiostat was built through FPGA board fabrication and assembly, similar to the Biospark potentiostat described in previous sections. Similarly, the system is connected to a PC via USB, and a custom Windows GUI program written in C# is used to control the system. Potentiostat settings for both channels (WE1 and WE2) such as voltage, pulse period and control settings such as current threshold can be adjusted in the program. Electrochemical data are saved in a custom binary format but can be exported into an Excel spreadsheet (.xlsx).

##### ***Custom algorithm for 'smart' control of enzymatic activity***

A simple algorithm was built to control the bio-electrochemical platform; the system diagram for the connecting both gene expression (by the local Biospark) and enzymatic activity (by the remote enzymatic checkpoint) is depicted in **Supplementary Figure 9**. After receiving the initiation message from the local Biospark system, the algorithm then commands the custom-built potentiostat (situated at a remote location) to run a pre-set program: apply -0.8 V on WE1 for 600 s, followed by 0 V on WE2 for 120s. It then compares the value of the output current to that of the user-defined current threshold: if the output current does not exceed the threshold, a message is sent to the local Biospark system to administer electroinduction immediately after the upcoming fluorescence measurement (for a user-defined duration or charge); otherwise, the algorithm sends a SMS verification message to alert the human users and seek permission to terminate the experiment. If 'Y' is received from the human users, a termination cue will be sent to the local Biospark system to initiate photobleaching immediately after the upcoming fluorescence measurement. If 'N' is received form the users, the algorithm then prompts the 'actuation checkpoint' to run the pre-set program once more.

### Supplementary Tables

***Supplementary Table 1. Strains and plasmids used in this study.***

| **Strains** | | |
|---|---|---|
| **Name** | **Genotype** | **Reference** |
| NEB10β | *E. coli Δ(ara-leu) 7697 araD139 fhuA ΔlacX74 galK16 galE15 e14- ϕ80dlacZΔM15 recA1 relA1 endA1 nupG rpsL* (Str$^R$) *rph spoT1 Δ(mrr-hsdRMS-mcrBC)* | New England Biolabs |
| DH5α-sfGFP | *E. coli* DH5α *attTn7::mTn7ΦsfGFP* | This study |
| NB101 | *E. coli* ZK126 Δ*rpoZ* | Bhokisham *et al.*[4] |
| JLD271 | *E. coli* K-12 *ΔlacX74 sdiA271::Cam* | Lindsay *et al.*[2] |
| BB170 | *V. harveyi luxN*::Tn5 | Surrette *et al.*[3] |
| **Plasmids** | | |
| **Name** | **Description** | **Reference** |
| pSC-S108gRNA | pSC101 ori, Kan$^r$, *soxR*, *soxRSp*, spacer 108, gRNA scaffold | Bhokisham *et al.*[4] |
| pOxy-LacZ-laa | pBR322 ori, Amp$^r$, proD pomoter, RBS 31, *oxyR*, *oxyRS* promoter, RBS 30, *lacZ* fused with LAA ssRA tag | Terrell *et al.*[5] |
| pSC-O108 | pSC101 ori, Kan$^r$, proD promoter, RBS31, *oxyR*, *oxySp*, spacer 108, gRNA scaffold | This study |
| pdCas9ω | ω was inserted into C termini of dCas9 in pdCas9-bacteria (Addgene plasmid # 44249), p15A ori, pLtetO-1, Cm$^r$ | Bhokisham *et al.*[4] |
| pMC-GFP | pWJ89 with pBR322 and Amp$^r$ instead of pSC101 ori and Kan$^r$ | Bhokisham *et al.*[4] |
| pMC-lasI-LAA | pMC-GFP *lasI* with the LAA ssRA tag instead of *gfpmut2* | This study |
| pLasR_S129T-GFPmut3 | pSB1A2 backbone, pTetR-LasR(S129T)-pLuxR-GFP | This study |
| pS1gRNA | *soxS* specific gRNA spacers S1 inserted into pgRNAbacteria (Addgene plasmid # 44251) with pBR322 ori, Amp$^r$, BBa_J23119 promoter | Bhokisham *et al.*[4] |
| pControlgRNA | Control spacer (from Bikard *et al.*[8]) in pgRNA-bacteria (Addgene plasmid # 44251) with pBR322 origin, Amp$^r$, J23119 promoter | Bhokisham *et al.*[4] |
| pLuxS1 | pS1gRNA with *luxS* specific gRNA spacer LuxS1 instead of S1 | This study |
| pSC-LuxS1 | pSC-O108 with *luxS* specific gRNA spacer LuxS1 instead of 108 | This study |
| pSC-sg108+LuxS1 | pSC101 ori, Kan$^r$, BBa_J23100 promoter, tracrRNA, b1002 terminator, proD promoter, RBS 31, *oxyR*, *oxyRS* promoter, DR, spacer 108, DR, spacer LuxS1, DR, b1006 terminator | This study |
| pOxy-sfGFP | pBR322 ori, Amp$^r$, proD pomoter, RBS 31, *oxyR*, *oxyRS* promoter, RBS 33, *sfGFP* | Li, Wang *et al.*[6] |
| pOxy-sfGFP-AAV | AAV ssRA tag inserted to *sfGFP* in pOxy-sfGFP | This study |

***Supplementary Table 2. Sequences of relevant genetic parts***

| **Name** | **Sequence (5'-3')** |
|---|---|
| proD promoter | AAAGTTAAACAAAATTATTTGTAGAGGGAAACCGTTGTGGTCTCCCTGAATAT ATTATACGAGCCTTATGCATGCCCGTAAAGTTATCCAGCAACCACTCATAGAC CTAGGGCAGCAGATAGGGACGACGTGGTGTTAGCTGTG |
| *oxyR* | ATGAATATTCGTGATCTTGAGTACCTGGTGGCATTGGCTGAACACCGCCATTTT CGGCGTGCGGCAGATTCCTGCCACGTTAGCCAGCCGACGCTTAGCGGGCAAAT TCGTAAGCTGGAAGATGAGCTGGGCGTGATGTTGCTGGAGCGGACCAGCCGT |

| | |
|---|---|
| | AAAGTGTTGTTCACCCAGGCGGGAATGCTGCTGGTGGATCAGGCGCGTACCGT GCTGCGTGAGGTGAAAGTCCTTAAAGAGATGGCAAGCCAGCAGGGCGAGACG ATGTCCGGACCGCTGCACATTGGTTTGATTCCCACAGTTGGACCGTACCTGCTA CCGCATATTATCCCTATGCTGCACCAGACCTTTCCAAAGCTGGAAATGTATCTG CATGAAGCACAGACCCACCAGTTACTGGCGCAACTGGACAGCGGCAAACTCG ATTGCGTGATCCTCGCGCTGGTGAAAGAGAGCGAAGCATTCATTGAAGTGCCG TTGTTTGATGAGCCAATGTTGCTGGCTATCTATGAAGATCACCCGTGGGCGAA CCGCGAATGCGTACCGATGGCCGATCTGGCAGGGGAAAAACTGCTGATGCTG GAAGATGGTCACTGTTTGCGCGATCAGGCAATGGGTTTCTGTTTTGAAGCCGG GGCGGATGAAGATACACACTTCCGCGCGACCAGCCTGGAAACTCTGCGCAAC ATGGTGGCGGCAGGTAGCGGGATCACTTTACTGCCAGCGCTGGCTGTGCCGCC GGAGCGCAAACGCGATGGGGTTGTTTATCTGCCGTGCATTAAGCCGGAACCAC GCCGCACTATTGGCCTGGTTTATCGTCCTGGCTCACCGCTGCGCAGCCGCTATG AGCAGCTGGCAGAGGCCATCCGCGCAAGAATGGATGGCCATTTCGATAAAGT TTTAAAACAGGCGGTTTAA |
| RBS BBa_B0031 | TCACACAGGAAACC |
| *oxyS* promoter | TATCCATCCTCCATCGCCACGATAGTTCATGGCGATAGGTAGAATAGCAATGA ACGATTATCCCTATCAAGCATTCTGACTGATAATTGCTCACA |
| Spacer 108 | GCGTGTTGTGGAAGATCCGGCCTGCAGCCA |
| gRNA scaffold | GTTTTAGAGCTAGAAATAGCAAGTTAAAATAAGGCTAGTCCGTTATCAACTTG AAAAAGTGGCACCGAGTCGGTGC |
| GFPmut2 | ATGAGTAAAGGAGAAGAACTTTTCACTGGAGTTGTCCCAATTCTTGTTGAATT AGATGGTGATGTTAATGGGCACAAATTTTCTGTCAGTGGAGAGGGTGAAGGTG ATGCAACATACGGAAAACTTACCCTTAAATTTATTTGCACTACTGGAAAACTA CCTGTTCCATGGCCAACACTTGTCACTACTTTCGCGTATGGTCTTCAATGCTTT GCGAGATACCCAGATCATATGAAACAGCATGACTTTTTCAAGAGTGCCATGCC CGAAGGTTATGTACAGGAAAGAACTATATTTTTCAAAGATGACGGGAACTACA AGACACGTGCTGAAGTCAAGTTTGAAGGTGATACCCTTGTTAATAGAATCGAG TTAAAAGGTATTGATTTTAAAGAAGATGGAAACATTCTTGGACACAAATTGGA ATACAACTATAACTCACACAATGTATACATCATGGCAGACAAACAAAAGAAT GGAATCAAAGTTAACTTCAAAATTAGACACAACATTGAAGATGGAAGCGTTC AACTAGCAGACCATTATCAACAAAATACTCCAATTGGCGATGGCCCTGTCCTT TTACCAGACAACCATTACCTGTCCACACAATCTGCCCTTTCGAAAGATCCCAA CGAAAAGAGAGACCACATGATCCTTCTTGAGTTTGTAACAGCTGCTGGGATTA CACATGGCATGGATGAACTATACAAA |
| *lasI*-LAA | ATGATCGTACAAATTGGTCGGCGCGAAGAGTTCGATAAAAAACTGCTGGGCG AGATGCACAAGTTGCGTGCTCAAGTGTTCAAGGAGCGCAAAGGCTGGGACGT TAGTGTCATCGACGAGATGGAAATCGATGGTTATGACGCACTCAGTCCTTATT ACATGTTGATCCAGGAAGATACTCCTGAAGCCCAGGTTTTCGGTTGCTGGCGA ATTCTCGATACCACTGGCCCCTACATGCTGAAGAACACCTTCCCGGAGCTTCT GCACGGCAAGGAAGCGCCTTGCTCGCCGCACATCTGGGAACTCAGCCGTTTCG CCATCAACTCTGGACAGAAAGGCTCGCTGGGCTTTTCCGACTGTACGCTGGAG GCGATGCGCGCGCTGGCCCGCTACAGCCTGCAGAACGACATCCAGACGCTGGT GACGGTAACCACCGTAGGCGTGGAGAAGATGATGATCCGTGCCGGCCTGGAC GTATCGCGCTTCGGTCCGCACCTGAAGATCGGCATCGAGCGCGCGGTGGCCTT GCGCATCGAACTCAATGCCAAGACCCAGATCGCGCTTTACGGGGGAGTGCTGG TGGAACAGCGACTGGCGGTTTCAGCAGCGAACGACGAAAATTACGCCCTTGC AGCGTGATAATAA |
| LasR (**S129T**) | ATGGCCTTGGTTGACGGTTTTCTTGAGCTGGAACGCTCAAGTGGAAAATTGGA GTGGAGCGCCATCCTGCAGAAGATGGCGAGCGACCTTGGATTCTCGAAGATCC TGTTCGGCCTGTTGCCTAAGGACAGCCAGGACTACGAGAACGCCTTCATCGTC GGCAACTACCCGGCCGCCTGGCGCGAGCATTACGACCGGGCTGGCTACGCGC GGGTCGACCCGACGGTCAGTCACTGTACCCAGAGCGTACTGCCGATTTTCTGG GAACCGTCCATCTACCAGACGCGAAAGCAGCACGAGTTCTTCGAGGAAGCCTC GGCCGCCGGCCTGGTGTATGGGCTGACCATGCCGCTGCATGGTGCTCGCGGCG |

| | |
|---|---|
| | AACTCGGCGCGCTG**ACC**CTCAGCGTGGAAGCGGAAAACCGGGCCGAGGCCAA<br>CCGTTTCATGGAGTCGGTCCTGCCGACCCTGTGGATGCTCAAGGACTACGCAC<br>TGCAGAGCGGTGCCGGACTGGCCTTCGAACATCCGGTCAGCAAACCGGTGGTT<br>CTGACCAGCCGGGAGAAGGAAGTGTTGCAGTGGTGCGCCATCGGCAAGACCA<br>GTTGGGAGATATCGGTTATCTGCAACTGCTCGGAAGCCAATGTGAACTTCCAT<br>ATGGGAAATATTCGGCGGAAGTTCGGTGTGACCTCCCGCCGCGTAGCGGCCAT<br>TATGGCCGTTAATTTGGGTCTTATTACTCTCTGA |
| Spacer LuxS1 | GGTATGATCGACTGTGAAGCTATCTAACAA |
| Spacer Control | TGAGACCAGTCTCGGAAGCTCAAAGGTCTC |
| crRNAs 108, LuxS1, and terminator BBa_b0016 | CTCGAGGTTTTAGAGCTATGCTGTTTTGAATGGTCCCAAAACGCGTGTTGTGG<br>AAGATCCGGCCTGCAGCCAGTTTTAGAGCTATGCTGTTTTGAATGGTCCCAAA<br>ACGGTATGATCGACTGTGAAGCTATCTAACAAGTTTTAGAGCTATGCTGTTTTG<br>AATGGTCCCAAAACAAAAAAAAACCCCGCCCCTGACAGGGCGGGGTTTTTTT<br>GGATCC |
| Promoter BBa_J23100, tracrRNA, and terminator BBa_b1002 | AAGCTTTTGACGGCTAGCTCAGTCCTAGGTACAGTGCTAGCGGAACCATTCAA<br>AACAGCATAGCAAGTTAAAATAAGGCTAGTCCGTTATCAACTTGAAAAAGTG<br>GCACCGAGTCGGTGCTTTTTTTCGCAAAAAACCCCGCTTCGGCGGGGTTTTTC<br>GCGACGTC |
| sfGFP-AAV | ATGAGCAAAGGAGAAGAACTTTTCACTGGAGTTGTCCCAATTCTTGTTGAATT<br>AGATGGTGATGTTAATGGGCACAAATTTTCTGTCCGTGGAGAGGGTGAAGGTG<br>ATGCTACAAACGGAAAACTCACCCTTAAATTTATTTGCACTACTGGAAAACTA<br>CCTGTTCCGTGGCCAACACTTGTCACTACTCTGACCTATGGTGTTCAATGCTTT<br>TCCCGTTATCCGGATCACATGAAACGGCATGACTTTTTCAAGAGTGCCATGCC<br>CGAAGGTTATGTACAGGAACGCACTATATCTTTCAAAGATGACGGGACCTACA<br>AGACGCGTGCTGAAGTCAAGTTTGAAGGTGATACCCTTGTTAATCGTATCGAG<br>TTAAAGGGTATTGATTTTAAAGAAGATGGAAACATTCTTGGACACAAACTCGA<br>GTACAACTTTAACTCACACAATGTATACATCACGGCAGACAAACAAAAGAAT<br>GGAATCAAAGCTAACTTCAAAATTCGCCACAACGTTGAAGATGGTTCCGTTCA<br>ACTAGCAGACCATTATCAACAAAATACTCCAATTGGCGATGGCCCTGTCCTTT<br>TACCAGACAACCATTACCTGTCGACACAATCTGTCCTTTCGAAAGATCCCAAC<br>GAAAAGCGTGACCACATGGTCCTTCTTGAGTTTGTAACTGCTGCTGGGATTAC<br>ACATGGCATGGATGAGCTCTACAAAGCAGCAAACGACGAAAACTACGCTGCT<br>GCTGTT |

***Supplementary Table 3. Primers used in this study.***

| Name | Sequence (5’-3’) |
|---|---|
| SW01 | TGTTTGACAGCTTATCATC |
| SW02 | GCGTCCGGCGTAGAGTCGTGTGAGCAATTATCAG |
| SW03 | CACACGAGCGTGTTGTGG |
| SW04 | GTAGAGAAAAAAGCACCGACTCGG |
| SW05 | GTCGGTGCTTTTTTCTCTACCTCTACGCCGGACGCATC |
| SW06 | TTCCACAACACGCTCGTGTGTCGTGTGAGCAATTATCAGTCAG |
| SW07 | GAAGCTATCTAACAAGTTTTAGAGCTAGAAATAGCAAGTTAAAATAAG |
| SW08 | ACAGTCGATCATACCTCGTGTGTCGTGTGAGCA |
| SW09 | CTAACGGATCCGCTTTTTTCTCTACCTCTACGCC |
| SW10 | CTAAGCTCGAGTGTGAGCAATTATCAGTCAGAATG |
| SW11 | GAAGCTATCTAACAAGTTTTAGAGCTAGAAATAGC |
| SW12 | ACAGTCGATCATACCACTAGTATTATACCTAGGAC |
| SW13 | GGATCCCATGGTACGCGTG |
| SW14 | CTAGATTTCTCCTCTTTAAAGGAATTCGC |

| | |
|---|---|
| SW15 | TTTAAAGAGGAGAAATCTAGATGATCGTACAAATTGGTCGG |
| SW16 | GCACGCGTACCATGGGATCCTTATTATCACGCTGCAAGGG |
| SW17 | TGATGCGGTAGTTTATCAC |
| SW18 | AACAGCAGCAGCGTAGTTTTCGTCGTTTGCTGCTTTGTAGAGCTCATCCAT |
| SW19 | GCGTGTTGTGGAAGATCCG |
| SW20 | CGACTCGGTGCCACTTTTTC |

#### Method References


1 Li, J. *et al.* Mediated Electrochemistry to Mimic Biology's Oxidative Assembly of Functional Matrices. *Advanced Functional Materials* **30**, 2001776, doi:https://doi.org/10.1002/adfm.202001776 (2020).

2 Lindsay, A. & Ahmer, B. M. Effect of sdiA on biosensors of N-acylhomoserine lactones. *J Bacteriol* **187**, 5054-5058, doi:10.1128/JB.187.14.5054-5058.2005 (2005).

3 Surette, M. G., Miller, M. B. & Bassler, B. L. Quorum sensing in *Escherichia coli, Salmonella typhimurium*, and *Vibrio harveyi*: a new family of genes responsible for autoinducer production. *Proc Natl Acad Sci U S A* **96**, 1639-1644, doi:10.1073/pnas.96.4.1639 (1999).

4 Bhokisham, N. *et al.* A redox-based electrogenetic CRISPR system to connect with and control biological information networks. *Nat Commun* **11**, 2427, doi:10.1038/s41467-020-16249-x (2020).

5 Terrell, J. L. *et al.* Bioelectronic control of a microbial community using surface-assembled electrogenetic cells to route signals. *Nat Nanotechnol* **16**, 688-697, doi:10.1038/s41565-021-00878-4 (2021).

6 Li, J. *et al.* Interactive Materials for Bidirectional Redox-Based Communication. *Advanced Materials* **33**, 2007758, doi:https://doi.org/10.1002/adma.202007758 (2021).

7 DeLisa, M. *et al.* Generic model control of induced protein expression in high cell density cultivation of Escherichia coli using on-line GFP-fusion monitoring. *Bioprocess and Biosystems Engineering* **24**, 83-91, doi:10.1007/s004490100229 (2001).

8 Bikard, D. *et al.* Programmable repression and activation of bacterial gene expression using an engineered CRISPR-Cas system. *Nucleic Acids Res* **41**, 7429-7437, doi:10.1093/nar/gkt520 (2013).
